# Disrupted developmental signaling induces novel transcriptional states

**DOI:** 10.1101/2024.09.05.610903

**Authors:** Aleena Patel, Vanessa Gonzalez, Triveni Menon, Stanislav Y. Shvartsman, Rebecca Burdine, Maria Avdeeva

**Author notes:** equal contribution (Aleena Patel, Maria Avdeeva).

## Abstract

Signaling pathways induce stereotyped transcriptional changes as stem cells progress into mature cell types during embryogenesis. Signaling perturbations are necessary to discover which genes are responsive or insensitive to pathway activity. However, gene regulation is additionally dependent on cell state-specific factors like chromatin modifications or transcription factor binding. Thus, transcriptional profiles need to be assayed in single cells to identify potentially multiple, distinct perturbation responses among heterogeneous cell states in an embryo. In perturbation studies, comparing heterogeneous transcriptional states among experimental conditions often requires samples to be collected over multiple independent experiments. Datasets produced in such complex experimental designs can be confounded by batch effects. We present Design-Aware Integration of Single Cell ExpEriments (DAISEE), a new algorithm that models perturbation responses in single-cell datasets with a complex experimental design. We demonstrate that DAISEE improves upon a previously available integrative non-negative matrix factorization framework, more efficiently separating perturbation responses from confounding variation. We use DAISEE to integrate newly collected single-cell RNA-sequencing datasets from 5-hour old zebrafish embryos expressing optimized photoswitchable MEK (psMEK), which globally activates the extracellular signal-regulated kinase (ERK), a signaling molecule involved in many cell specification events. psMEK drives some cells that are normally not exposed to ERK signals towards other wild type states and induces novel states expressing a mixture of transcriptional programs, including precociously activated endothelial genes. ERK signaling is therefore capable of introducing profoundly new gene expression states in developing embryos.

**Significance Statement:** Signaling perturbations produce heterogeneous transcriptional responses that must be measured at the single-cell level. Data integration techniques allow us to model these responses which, however, can be confounded by batch effects. We present a computational tool (DAISEE) for extracting the common and perturbation-specific features of single-cell datasets representing multiple experimental conditions while achieving efficient batch effect correction. DAISEE outperforms its predecessor and will enable accurate analysis of a broad range of single-cell datasets. DAISEE applied to new single-cell RNA sequencing data from zebrafish embryos shows that gain-of-function signaling perturbations can induce novel states. Our analysis suggests that a wild type endothelial cell-specification program can be activated in abnormal developmental contexts when the extracellular signal-regulated kinase (ERK) pathway is deregulated.

## Introduction

During metazoan embryogenesis, pluripotent cells acquire specific identities via systematic changes in gene expression, in large part coordinated by the stereotyped activation of cell-cell signaling pathways. A signaling perturbation can generate diverse and context-dependent gene expression responses that are challenging to predict *a priori* and can only be fully described using single-cell measurements. Genomic technologies can provide unbiased, comprehensive readouts of individual cell states. Comparing these readouts upon perturbation to wild type embryos may reveal specific signal-driven changes in gene regulatory programs that derail normal cell specification and cause developmental disease. Here, we develop a computational tool for integrating single-cell perturbation experiments and use it to quantitatively characterize single-cell transcriptional responses to global signaling activation in early zebrafish embryos.

In response to changes in signaling, cell states in an embryo can become redistributed, undergo a minor perturbation, or entirely different states unseen in the wild type embryo can be realized. To obtain a comprehensive description of perturbation responses in these terms, it is necessary to collect large datasets of high-dimensional single-cell measurements corresponding to multiple conditions. To increase statistical power, data from many cells are often gathered in several independent experiments, and the responses to perturbation can be confounded by sources of variation other than the perturbation itself, such as batch effects due to unavoidable technical differences in experimental protocol or sequencing procedure. Computational integration techniques can provide a unified view of all samples by embedding the measured cell states into a common phenotypic space and modeling the perturbation responses of interest, potentially separating them from confounding variation (1–5). Our tool is designed to improve batch effect correction in a leading family of perturbation modeling techniques by incorporating information about the experimental design into the algorithm.

We perform extensive benchmarking to demonstrate that our refined approach, Design-Aware Integration of Single Cell ExpEriments (DAISEE), more efficiently separates perturbation effects from confounding variation introduced by batch effects than its predecessor, Linked Inference of Genomic Experimental Relationships (LIGER) (1), in simulated, existing, and new data. DAISEE is a non-negative matrix factorization (NMF) technique, which extracts state signatures as gene modules, or factors, and has been shown to be an effective approach for interpreting embryonic data that lie on a continuum, with every cell expressing a combination of gene modules (6). Our algorithm applies broadly to any perturbation experiment that produces high-dimensional, single-cell measurements and requires a known source of confounding variation to be corrected for.

With DAISEE, we integrate newly collected single-cell RNA sequencing (scRNA-seq) data from 5 hour old zebrafish embryos experiencing constitutive activation of the conserved extracellular signal-regulated kinase (ERK) pathway, which coordinates cell fate specification widely in vertebrate embryogenesis (7–9). We sustain ERK activity without relying on the endogenous signal transduction cascade by using an optogenetic tool, photoswitchable MEK (10). Our study is the first to assay single-cell responses to overactive signaling in an embryo, which we hypothesized would expose many different cells to abnormal differentiation cues and potentially reveal new gene expression profiles.

We indeed find novel transcriptional states driven by multiple DAISEE factors, including one that resembles a transcriptional signature of endothelial fate appearing much later in development. This work illustrates remarkable plasticity of cells in the zebrafish embryo, and highlights the need to push signaling activity beyond the endogenous spatiotemporal limits to discover gene responses that are unrealized in wild type embryos, but potentially play important roles in driving disease phenotypes. We anticipate that DAISEE will be useful in continuing to study more sophisticated signaling perturbations that can be made with a variety of optogenetic signaling tools now available for multiple developmental pathways (10–13).

### Single-cell RNA sequencing of optogenetically ERK-activated zebrafish embryos

We performed scRNA-seq on dissociated cells from zebrafish embryos (Fig. 1a). We assayed the 50% epiboly stage, when the thinning and spreading of the blastoderm over the yolk, in a process called epiboly, reaches the halfway point. At this stage, a handful of signaling cues, including the Fibroblast Growth Factor (FGF), Nodal, Bone Morphogenetic Protein (BMP), and Wnt pathways, are subject to strict spatiotemporal regulation and induce non-uniform expression of key cell fate determining genes (8, 14). FGFs are ligands that bind to FGF receptors (FGFRs) and then trigger a phosphorylation cascade from the kinases RAF to MEK to ERK (Fig. 1b). Phosphorylated ERK is the active effector molecule of the pathway, and translocates to the nucleus to interact with transcription factors and regulate gene expression. We changed activity of the FGF/ERK pathway, which is normally restricted to the dorsal side of the embryo before epiboly starts, then shifts to the blastoderm-yolk margin during epiboly (15–18). The FGF-driven ERK activity contributes to the specification of dorsal versus ventral cell fates and germ layer (mesoderm, endoderm, and ectoderm) fates (19–21).

**Figure 1.**
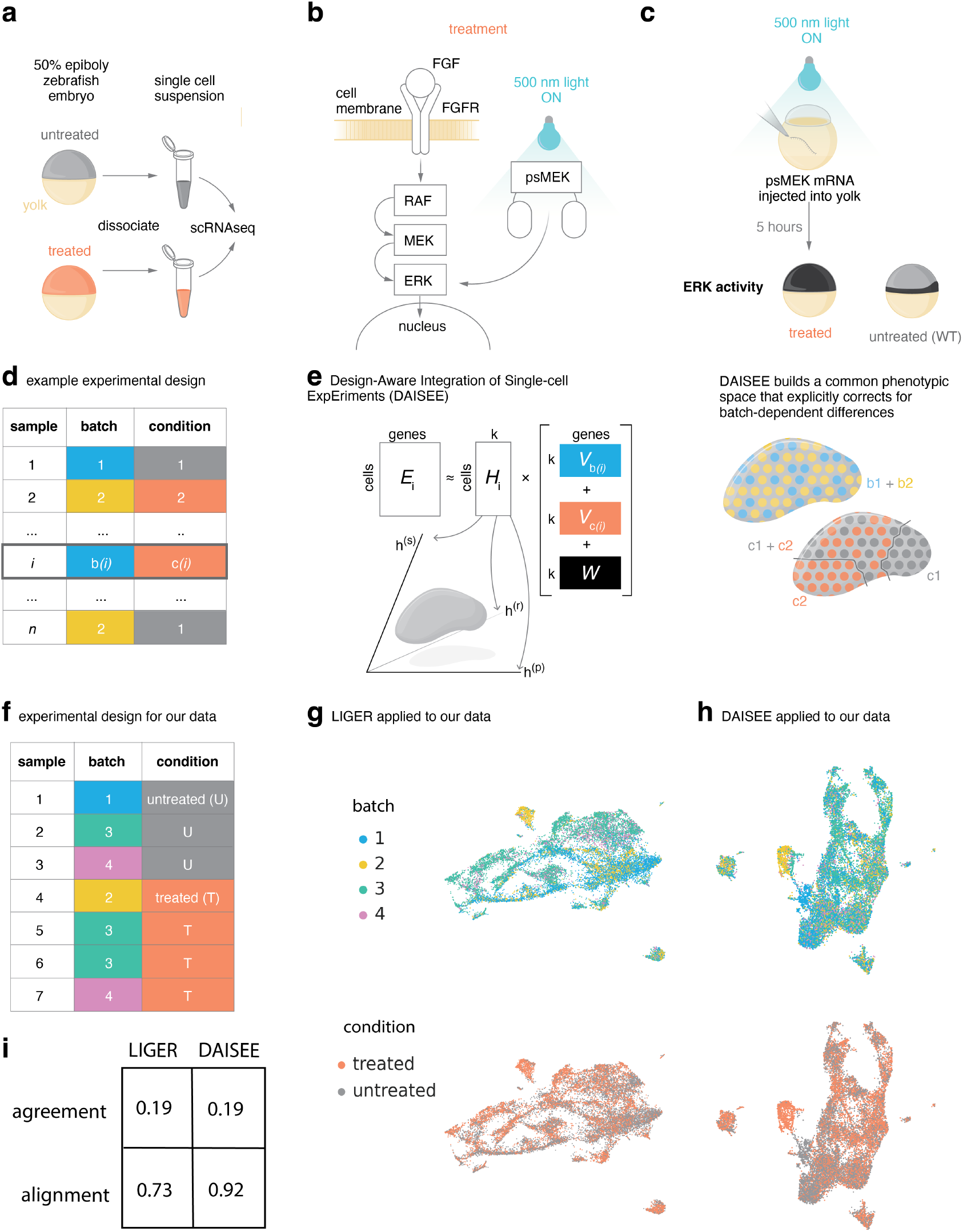
DAISEE provides efficient integration of perturbation experiments. a) Our data come from untreated and treated zebrafish embryos at the 50% epiboly stage. Embryos were dissociated in preparation for single-cell RNA sequencing. b) The treatment affects the ERK pathway. In the wild type embryo, fibroblast growth factor (FGF) ligands bind to FGF receptors (FGFRs) to trigger a phosphorylation cascade from the kinases RAF to MEK to ERK. In treated embryos, an optogenetic tool (psMEK) bypasses the signal transduction cascade and directly activates ERK. 500 nm light uncages photodimerizable domains in psMEK that permit its activation of ERK. Active ERK translocates to the nucleus to regulate gene expression. c) psMEK was expressed by injecting mRNA encoding psMEK into the yolk of the 1-cell stage embryo. Injected embryos were then exposed to 500 nm light for 5 hours while they developed. This treatment activates ERK throughout the blastoderm, whereas in untreated wild type embryos, ERK activity (black) is restricted to the margin. d) A table describing an experimental design with *n* single-cell samples collected over multiple batches and corresponding to different conditions. Sample *i* corresponds to batch *b*(*i*) and condition *c*(*i*). e) A schematic of Design-Aware Integration of Single-cell ExpEriments (DAISEE) algorithm. DAISEE jointly decomposes single-cell datasets, *E*_*i*_, into products of non-negative matrices (of lower rank *k*). *H*_*i*_: factor scores (loadings), *W* : common factors, *V*_*b*(*i*)_: batch-specific factors, *V*_*c*(*i*)_: condition-specific factors. This decomposition allows us to explicitly correct for batch dependent differences for reliable downstream interpretation of condition-dependent differences. f) The experimental design table for newly collected data. g) UMAPs showing batch-dependent and condition-dependent differences for the newly collected zebrafish data integrated using LIGER. h) Same as g) for DAISEE. i) Sample agreement and batch alignment for DAISEE vs LIGER on zebrafish embryonic data.

We implemented an optogenetic version of mitogen-activated protein kinase kinase (MEK) (photoswitchable MEK, psMEK) to add an orthogonal source of direct, FGF-independent ERK activation and perturb the signaling landscape of the embryo (22) (Fig. 1b). We had previously optimized psMEK using several gain-of-function mutations that lock MEK in its active conformation without phosphorylation by RAF (10). psMEK contains photodimerizable domains that harness this ligand-independent constitutive activity.

We injected mRNA for optimized psMEK into embryos at the 1-cell stage, and let them develop for 5 hours while exposed to the psMEK-activating wavelength of light (500 nm) (Fig. 1c). At the 50% epiboly stage, active ERK, indicated by immunofluorescence staining of dually phosphorylated ERK, is only found at the blastoderm-yolk margin and not in the remainder of the blastoderm (animal pole) (10, 23). In previous work, we showed ERK activation throughout the blastoderm, outside of the margin, using similar immunofluorescence stainings in 50% epiboly embryos treated with optimized psMEK exactly as we have done in this study (10). Our psMEK treated embryos therefore experience strong, ectopic ERK activation, which could result in cells in the animal pole assuming states normally driven by FGF signals in other parts of the embryo, or new states that emerge as a result of treatment-induced new combinations of ERK signals with other active signals in the embryo.

### DAISEE provides efficient integration of perturbation experiments

To obtain a comprehensive description of the effects of a perturbation on cell states, including those on rare cell types, it is necessary to collect single-cell data from thousands of cells. To this end, new data for the experimental conditions of interest are often acquired in several batches, or data are integrated from independent previous experiments (example experimental design in Fig. 1d). However, batch effects confound the signal of interest, and must be accounted for to interpret condition-dependent effects (Fig. 1e).

To efficiently integrate our dataset and separate the effects of perturbation from batch effects, we developed DAISEE, an algorithm that is broadly applicable to any perturbation study with a complex experimental design (Fig. 1d). DAISEE provides a joint embedding of all the cells into a common low-dimensional space, correcting for batch effects and aligning cells before and after perturbation. DAISEE is based on a previously available integrative NMF (iNMF) framework (1, 24) and is a simple and convenient tool for perturbation experiment analysis and interpretation. To integrate the data, iNMF extracts gene signatures (factors) that can usually be interpreted as a biological signal such as a transcriptome of a particular cell state or a transcriptomic signature of a particular developmental program. DAISEE models the perturbation affecting every factor, simultaneously extracting and correcting the factor-specific batch effects. Ultimately, for each factor, every cell gets assigned an expression score which not only serves to construct the joint embedding but can also be used for quantitative interpretation of cell states.

More precisely, DAISEE achieves efficient batch correction and perturbation modeling by explicitly incorporating the known experimental design into an iNMF model and separating the effects of a confounding factor from an effect of interest like variation due to perturbation. iNMF simultaneously decomposes the normalized count matrices from the single-cell datasets into products of lower rank non-negative matrices (Fig. 1e). While some gene signatures are shared between all datasets (common factors, *W*, Fig. 1e), there are other sources of biological and technical variability that are specific to subsets of the data and are captured by sample-specific factors *V*_*c*(*i*)_ and *V*_*b*(*i*)_. While batch-specific factors *V*_*b*_ allow for batch effect correction, condition-specific factors *V*_*c*_ can be further interpreted to analyze the effects of perturbation.

We applied DAISEE to our experimental design which included 7 samples representing 2 conditions and 4 batches (24,957 cells, Fig. 1f) after extensively benchmarking DAISEE’s performance against its iNMF predecessor, LIGER (1) (Supp. Figs.1,2). LIGER is a competitive method of batch effect correction for scRNA-seq data and has been shown to outperform other methods in a scATAC-seq data integration task in an extensive benchmarking study (25). However, when integrating perturbation conditions, LIGER does not directly incorporate the batch effects into its objective function and corrects for batch effects only implicitly through a joint clustering and quantile normalization procedure applied to the factor scores post-factorization. We observed that providing DAISEE with explicit information on experimental batches resulted in better batch effect correction and retained the ability to integrate conditions in our experiment (Fig. 1g,h).

Batch effect correction efficiency was measured using 2 previously introduced metrics, sample agreement and batch alignment (1, 26). Sample agreement measures the amount of distortion that the samples undergo during embedding into the common phenotypic space. At the same time, the batch alignment metric reflects how well-mixed the batches are in local neighborhoods on average, with perfectly mixed batches getting a score of 1. DAISEE, while preserving the level of agreement between samples, improves alignment as compared to LIGER (Fig. 1i).

To test the performance of DAISEE on the batch effect correction task for different perturbation strengths over a range of regularization parameters, we simulated single-cell transcriptomic data from the iNMF framework, using a simple experimental design with 2 treatment conditions collected over 2 different batches (Methods, Supp. Fig. 1a,b). We found that on simulated data DAISEE significantly outperforms LIGER for both agreement and alignment metrics, with increase in performance becoming larger as we increased the strength of the batch effect (Supp. Fig. 1c,d). To further increase the complexity of the integration task, we additionally benchmarked DAISEE on two different previously available atlases of human and mouse immune cells and cells of the mouse embryo (25). We explored a range of regularization strengths for batch-specific components and uncovered that significant alignment improvement over LIGER can be achieved without loss of agreement on both datasets (Supp. Fig. 2a,b).

Collectively, these results suggest that DAISEE allows for an improved integration of single-cell datasets. Thus, we apply it to study psMEK-driven perturbation effects in early zebrafish embryos.

### Annotation of cell states embedded in the DAISEE space

To analyze our zebrafish embryonic data with DAISEE, we performed additional tuning of the parameters (Methods). We retained *k* = 30 components for the analysis based on the Kullback-Leibler (KL) divergence heuristic as suggested in (1) (Supp. Fig. 3a). Additionally, we tuned the regularization parameter of DAISEE controlling the magnitude of the batch-specific factors (Supp. Fig. 3b). Furthermore, after applying DAISEE, we identified 9 factors that were highly batch-specific and manually filtered them out which resulted in a small increase of alignment at the expense of a small decrease of agreement (Methods, Supp. Fig. 3d). We proceeded with the resulting 21 DAISEE factors and used them to construct an ambient space with embedded cell states from our data (Fig. 2a). We scaled iNMF factors and partitioned the cells into discrete clusters of states based on the factor with the highest score (Fig. 2b).

**Figure 2.**
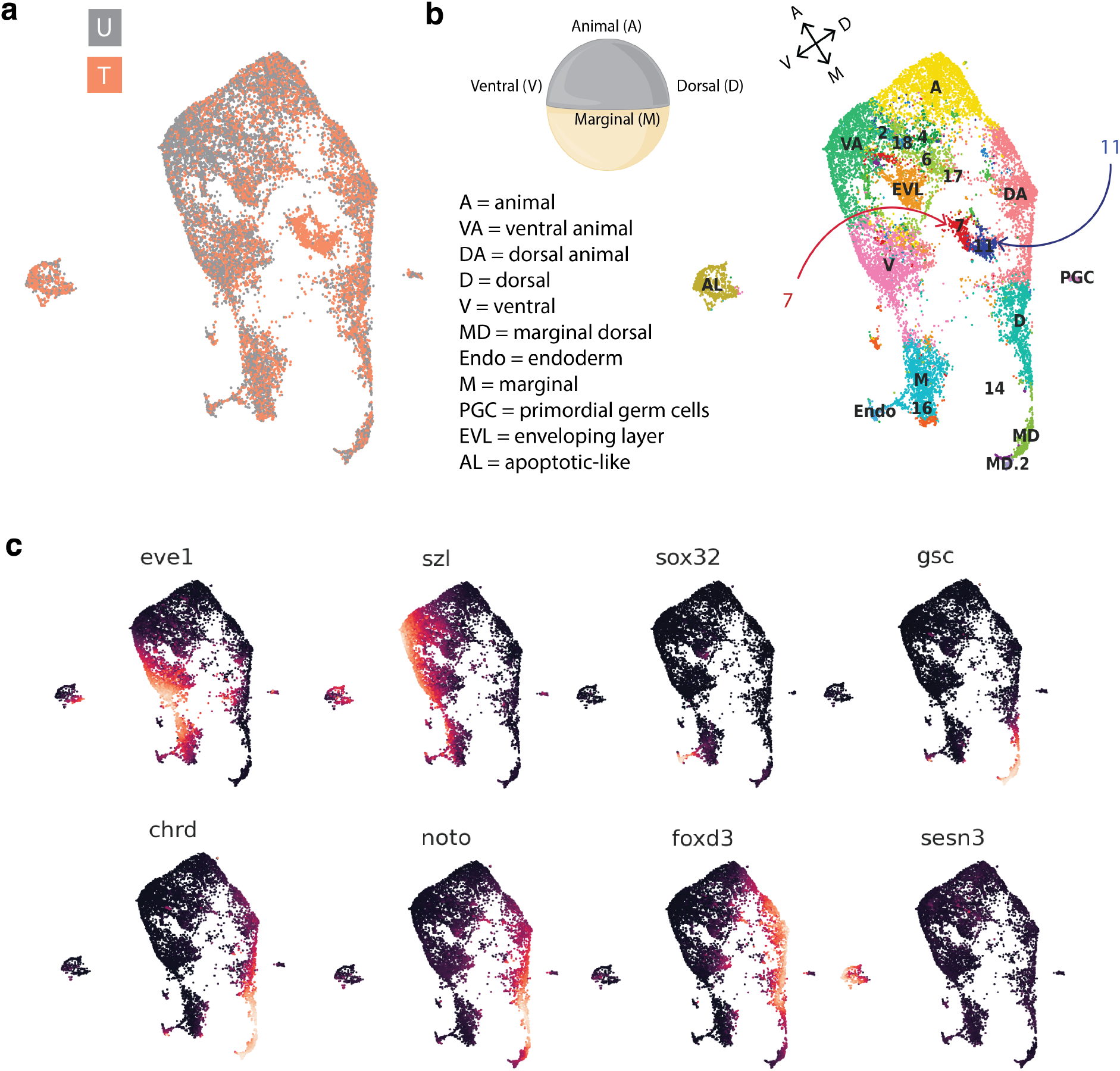
Annotation of cell states in psMEK treated perturbation experiments. a) UMAP of zebrafish embryonic data integrated with DAISEE. Orange corresponds to data from the treated condition (T) and grey corresponds to data from the untreated condition (U). b) Axes extending from the dorsal (D) side to the ventral (V) side and from the margin (M) to the animal pole (A) of 50% epiboly zebrafish embryo define a coordinate system on its surface. DAISEE clusters are shown; each cluster corresponds to a dominating factor. Clusters were annotated using previous publications; several clusters are annotated using the A-M, V-D coordinate system of the 50% epiboly zebrafish embryo. c) UMAPs showing expression of select gene markers in the untreated data (imputed over 50 nearest neighbors).

We turned to published data to annotate the factors and corresponding cell states captured in our experiments. We compared DAISEE factors to annotated NMF factors extracted from previously collected scRNA-seq data from wild type 50% epiboly zebrafish embryos reported in Farrell et. al. (6) (Methods, Supp. Fig. 3c). We identified clusters based on all of the published and annotated spatial factors (labeled by their ventral (V) versus dorsal (D) or animal (A) versus marginal (M) expression patterns), as well as the enveloping layer cells (EVL), the previously reported apoptotic-like (AL) cells, and the primordial germ cell (PGC) cells (6, 27). We additionally annotated one of the DAISEE factors and associated cluster as endoderm (Endo) due to its high expression of sox17 and sox32 in its common component *W*. Several states could not be annotated, which we identified with the number of the factor with the highest expression. This annotation allowed us to interpret quantitative effects of treatment in terms of known cell types.

We provide the gene expression profiles of several representative markers in the DAISEE integration in Fig. 2c. To highlight a few top gene markers, *eve1* (factor V) is a transcription factor that promotes ventral fates and is under control of ventralizing cues antagonized by dorsalizing factors like *chrd* (factor D) produced on the opposite side of the embryo (28, 29). The dorsal shield, equivalent to the Spemann organizer, is marked by *gsc* (factor MD) (30, 31). *sesn3* (factor AL) is a gene marker for the rare apoptotic-like cell type first reported in (27). A dotplot that summarizes per-cluster expression of top 10 genes of the common component of each factor can be found in Supp. Fig. 4. In sum, DAISEE integration produced a map of cell states in which mRNA signals for genes with known biological function are detected in expected patterns, and DAISEE factors can be used to partition the map into interpretable regions.

### psMEK treatment redistributes states in the DAISEE space

Since FGF plays a key role in cell fate determination, we posited that out-of-bounds ERK signaling in psMEK-treated embryos could expand some states at the expense of others. We sought to identify the cell states that were over-or underrepresented in the perturbed condition when compared to control. Towards this end, we applied MILO to test for differential abundance of conditions in local neighborhoods of the integrated dataset (32). MILO builds a k-nearest neighbors (kNN) graph of all the cells using a sampling procedure to define overlapping neighborhoods of the graph that can then be tested for differential abundance (Methods). By studying how cell states were redistributed by treatment, we could identify which transcriptional states were susceptible to changes in signaling conditions.

We identified multiple neighborhoods that were differentially abundant between treatment conditions at a spatial false discovery rate threshold of 5% calculated by MILO (Fig. 3a, Supp. Fig. 5a). After mapping neighborhoods back to cluster annotations through majority voting, we discovered differential abundance within several regions of the manifold (Fig. 3a). The most striking trends in the redistribution profile overall pointed to a large depletion in the VA region and enrichment in the DA region, and enrichment of the AL and unnamed 7 and 11 states. A more detailed substructure of the redistribution within neighborhood clusters, evident in the DAISEE framework, showed that there is a correlation between a cell’s DA or VA factor expression and how strongly psMEK treatment enriches or depletes the DA or VA state respectively (Supp. Fig. 5b). This result is consistent with psMEK treatment deregulating the cell fate specification programs that establish spatially variable cell states along the emerging dorsoventral axis. In summary, psMEK treatment caused changes in specification of the VA state that led to its loss, while also promoting the overspecification of other states.

**Figure 3.**
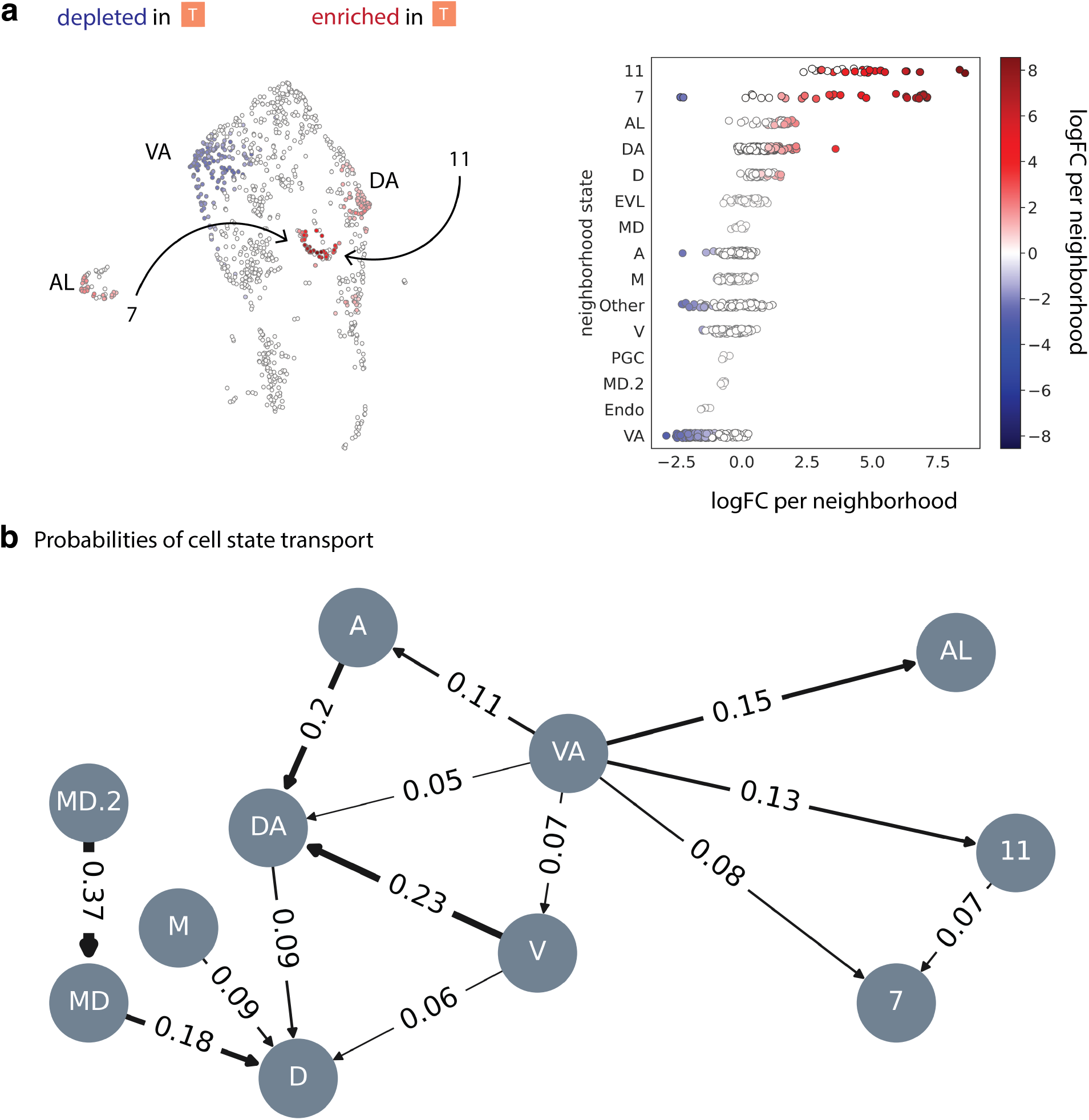
psMEK treatment redistributes cell states. a) Differential abundance testing (MILO) reports neighborhood states that are enriched (red) or depleted (blue) in the psMEK treated condition. Left: Per-neighborhood results are summarized in the UMAP. Right: Every dot corresponds to one of the overlapping neighborhoods formed by MILO. *x*-axis: log-foldchange of abundance (logFC) per neighborhood of treated vs untreated conditions, *y*-axis: DAISEE cluster annotation transferred to the neighborhood, color: logFC per neighborhood, with only neighborhoods below 5% Spatial FDR cutoff shown in color. b) Transport probabilities between spatially variable (VA, A, DA, V, MD, D, M, MD.2) and clusters 7, 11 and AL. The transition probabilities are derived from the optimal transport map 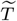. Every edge is marked with the corresponding transition probability and its width is proportional to it.

### Optimal transport underlying cell state redistribution describes likely paths of signal-driven reprogramming

We next sought to build a map of cell state reassignments, driven by the changes in gene expression caused by psMEK treatment, that would result in the redistribution detected by MILO. Optimal transport, which is widely studied in mathematics and applies to many problems of resource allocation, generally seeks to map one probability distribution into another one while minimizing the overall cost of the mapping for a predefined cost function. Optimal transport algorithms have been applied to scRNA-seq data to extract trajectories or study perturbation responses (4, 5, 33, 34). In a toy example, the probability distributions in question represent a pile of sand and a sand castle with the cost function defined to be the cost of transport from location *x* in the pile to location *y* in the castle (Supp. Fig. 6a). In our datasets, the probability distributions we wish to map are the distributions of cell states (over MILO neighborhoods) in the untreated and treated conditions (Supp. Fig. 6c). We chose Kantorovich relaxation of the optimal transport problem where, for every untreated cell state *x*, a transportation plan is stochastic and describes a probability distribution of treated cell states *y* to which *x* is mapped as a result of perturbation.

The resulting map of transport probabilities (Fig. 3b) reports a reassignment plan that includes shifts in cell state that were not apparent in the global redistribution. In addition to direct reassignment of the VA state to other enriched states, which would result in depletion of VA cells and enrichment of other cells as we have observed, there are several transport paths that include multiple steps through intervening states (see multiple arrows from the VA state to other spatially variable states in Fig. 3b). Such gradual transportation is not observed in the transport of the VA state towards the enriched non-spatially variable states AL, 7, and 11, and could indicate different psMEK-driven transcriptional changes in the enrichment of spatially-variable states versus the other states.

The step-wise transport of cells through wild type spatially variable states (VA, A, DA, V, MD, D, M, MD.2) could reflect regulation of a differentiation cue that responds in a graded manner to ERK perturbation. We posit that this behavior is the attenuation of the BMP signaling gradient that acts as a master regulator of dorsoventral fates across vertebrates (35). In psMEK treated embryos, BMP signaling is indeed attenuated at multiple levels of the signaling cascade. Prior measurements of p-Smad1/5/8 phosphorylation, the active effector molecule of the BMP pathway, in psMEK treated embryos showed loss of signaling on the ventral side of the embryo (10). This study’s scRNA-seq found upregulation of *chrd*, a well-known antagonist of BMP ligand-receptor binding and downregulation of *bmp2b*, which encodes a ligand that activates the BMP pathway (Supp. Fig. 6d). Loss of BMP signaling in ventral cells would fail to specify ventral fates and instead permit over-representation of signaling conditions that promote dorsal fates. The systematic shift towards more dorsal states points to the early embryo undergoing dorsalization.

The reassignment of VA cells directly to the non spatially variable states (AL, 7, 11) remained unclear. Apoptotic-like (AL) cells are a rare cell type found primarily in the animal pole (6). The cell states found in cluster 7 and 11 are foreign to the wild type 50% zebrafish embryo, yet are not artifacts of confounding factors and are present in all replicates of psMEK treatment. We note enrichment of these states was not a result of embryos exposed to heat or light during optogenetic treatment (Supp. Fig. 7). We posited that there were new gene expression behaviors in embryos responding to psMEK treatment that cannot be explained by well-known transcriptional signatures or gene-regulatory phenomena in the wild type 50% epiboly zebrafish embryo.

### New endothelial and stress-like factors shape the DAISEE space

We turned to the gene markers for the common component of factors 7 and 11 reported in the *W* term of DAISEE to search for identities of the significantly enriched new states. Factors 7 and 11 were both expressed in differentially abundant cells, which suggests that these transcriptional programs simultaneously drive the emergence of the new states (Fig. 4a). By visual inspection, we noticed several genes (several heat shock protein *hsp* genes, *ubb, dusp5*, see Fig. 4b) in the list of top 100 markers for factor 7 that matched markers of a transcriptional program found in a “stress-like” cancerous state (36). Cells in this state were discovered in a scRNA-seq study of melanoma tumors induced by expressing the human oncogene BRAFV600E, which affects the ERK pathway, in zebrafish. One of the constitutively activating mutations (E203K) in psMEK is also associated with cancer (37, 38). We concluded that we had triggered a stress response, and labeled factor 7 stress-like (SL).

**Figure 4.**
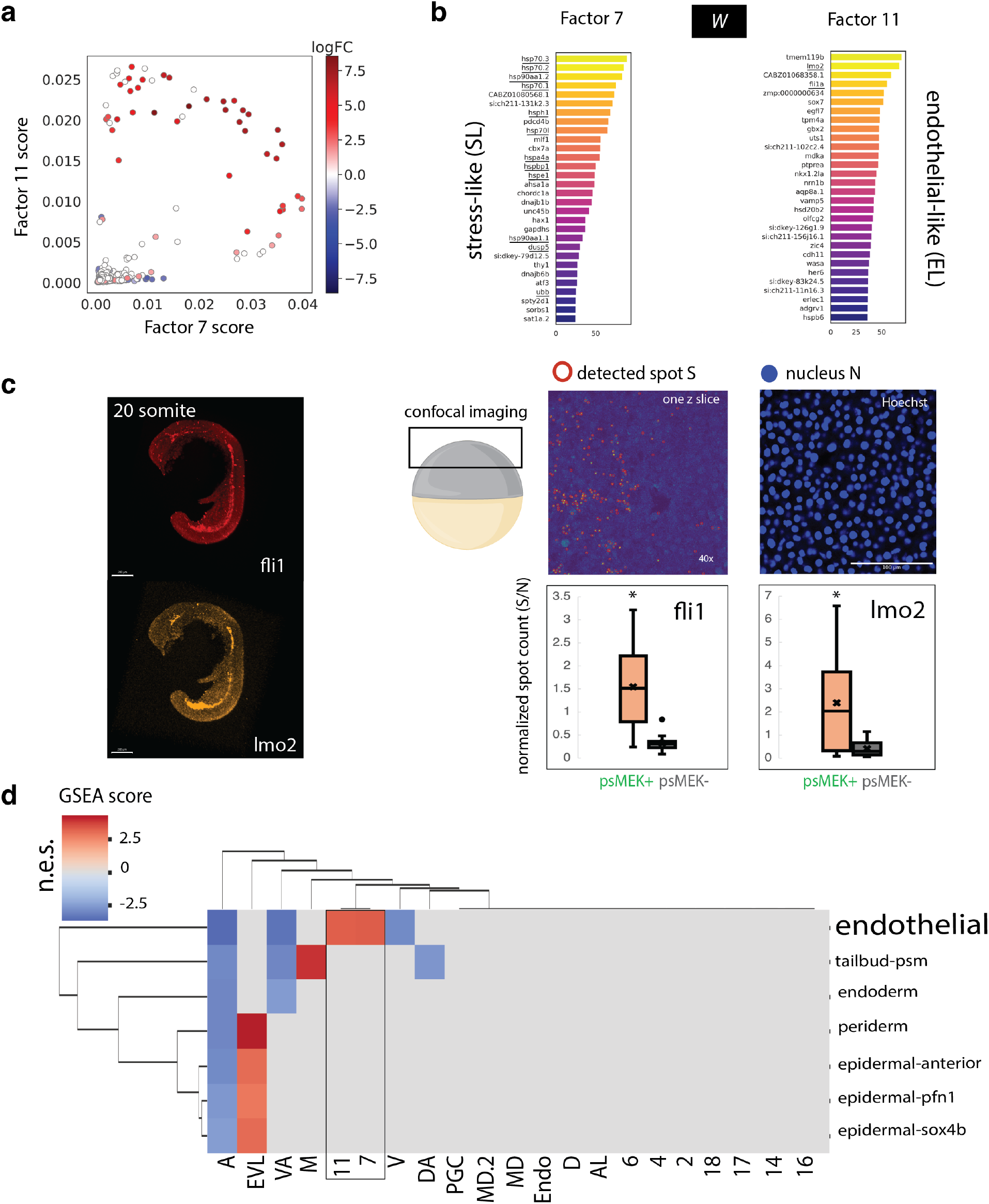
psMEK treatment-specific DAISEE factors are endothelial-like and stress-like. a) Mixing of factors 7 and 11. *x*-axis: per-neighborhood factor 7 score, *y*-axis: same for factor 11. Per-neighborhood scores were calculated by averaging cell scores over the neighborhood. Neighborhoods colored by their logFC in treated-vs-untreated cell abundance (as in Fig. 3a). b) Gene markers for the common (shared, *W*) component of factors 7 (stress-like) and 11 (endothelial-like). Key genes identifying the factor are underlined. c) Left: RNA Hybridization Chain Reaction *in situ* images for *fli1* and *lmo2* in wild type 20 somite zebrafish embryos. Scale bar = 200*µ*m. Right: The same probes were used to label *fli1* and *lmo2* in 50% epiboly stage embryos imaged in the animal pole at 40x. All embryos were co-stained with Hoechst. In each z slice, the RNA spots for *fli1* and *lmo2* were detected and counted, as were number of nuclei. The number of spots divided by the number of nuclei (S/N) is reported. In the injected embryos, only cells expressing psMEK (psMEK+) were considered. All cells in the uninjected (psMEK-) images were measured. For the *fli1* HCR in situ: 14 images from 3 injected embryos and 17 images from 4 uninjected embryos were analyzed. For the *lmo2* HCR in situ: 22 images from 5 injected embryos and 17 images from 4 uninjected embryos were analyzed. A t-test indicates statistically significant (*p<* .05) differences (*) between the psMEK+ and psMEK-data. d) Normalized enrichment score after performing GSEA. 18 h.p.f. endothelial genes were highly differentially expressed in the 7 and 11 clusters of cell states (boxed). FDR q-value *<* .001 cutoff was applied.

The genes *lmo2* and *fli1* found in factor 11 (Fig. 4b) encode transcription factors that play key roles in the normal specification of endothelial cells (39, 40). While endothelial precursors are specified from the ventral mesoderm of the early embryo, *lmo2* and *fli1* are not expected to be highly expressed throughout the early embryo, and rather are expressed abundantly in the vasculature of the much older somite-stage embryo (41). We investigated their expression using gene-specific probes that we could label and image at multiple stages of embryogenesis. We used HCR RNA-FISH probes to label *lmo2* and *fli1* transcripts in the emerging vasculature of somite-stage embryos (Fig. 4c). We then used these same probes to label *lmo2* and *fli1* transcripts in the wild type 50% epiboly embryo, and indeed found little expression. However, in psMEK-treated 50% epiboly embryos stained and imaged in the same experiment we found abundant labeling of *lmo2* and *fli1* transcripts (Fig. 4c). Therefore, in psMEK-activated conditions, *lmo2* and *fli1* appear to be upregulated in parts of the embryo that normally do not express these genes.

We explored the hypothesis that factor 11 represents a gene signature of a precocious endothelial fate by taking advantage of a published comprehensive transcriptomic atlas of zebrafish development. We performed differential expression analysis between the cell states identified in our datasets to search for enrichment of gene sets from every stage of embryogenesis reported in (42). This atlas contains 198 gene sets of markers of cell states from zebrafish aged 4 h.p.f. to 24 h.p.f. Gene set enrichment analysis (43) showed that the new clusters 7 and 11 significantly overexpress a gene set belonging to a cell state found at 18 h.p.f. and corresponding to an endothelial fate (Fig. 4d, Supp. Fig. 8a). Several gene markers from the 18 h.p.f. endothelial transcriptional program were exclusively expressed in the new clusters 7 and 11 (Supp. Fig. 8b). Thus, we named factor 11 endothelial-like (EL).

The new SL and EL psMEK-specific factors represent highly abnormal transcriptional signatures expressed in the treated embryos. The EL and SL factors expand the set of possible gene expression states, represented in our framework as an ambient space built by DAISEE.

### Quantitative changes in DAISEE factors describe perturbation responses

With new factors annotated, we could explore factor score changes during the cell state reassignment to build a quantitative picture of the perturbation responses. In the zebrafish embryo, one of the first cell fate decisions requires signals derived from the dorsal organizer, together with BMP signals, to define dorsal versus ventral cell types. It has long been proposed that FGF ligands are a part of these dorsalizing cues (19), but until now characterization of signal-driven dorsalization has been limited by the number of genes that could be labeled. DAISEE provides a quantitative description of complex changes in gene expression during dorsoventral reprogramming at the whole transcriptome level. We found a systematic loss of expression of the VA factor across states, accompanied by gain in expression of the DA factor (Fig. 5a). The MD factors expressing *gsc* were not as upregulated, thus, psMEK treatment appears to have promoted shifts towards dorsal states without expanding the progenitors of the dorsal organizer itself.

**Figure 5.**
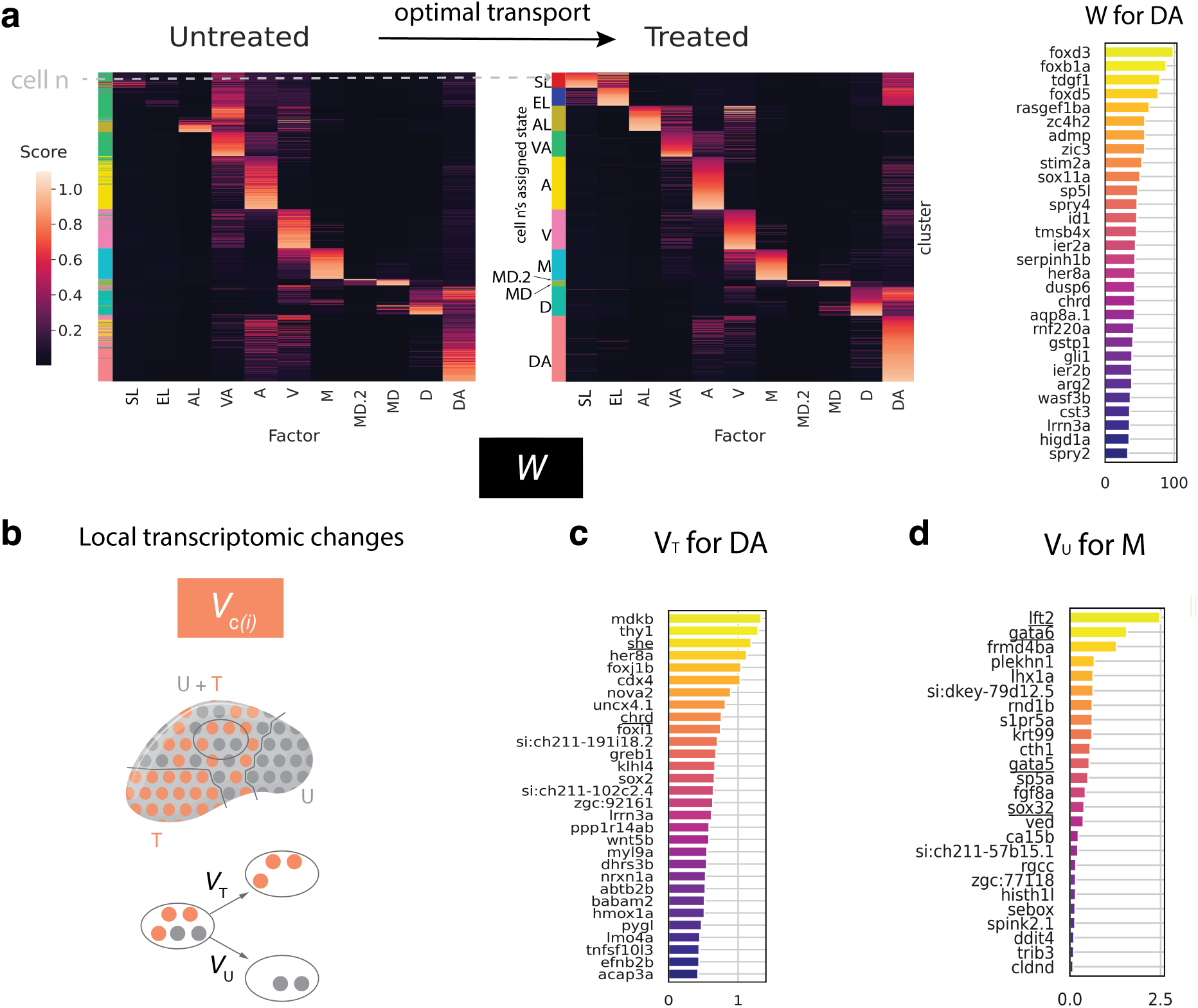
Quantitative changes in DAISEE factors describe perturbation responses. a) Heatmaps: factor scores before and after treatment are shown, with the rows on the left and right in direct correspondence reflecting the reprogramming changes derived from the optimal transport analysis. More precisely, in the heatmap on the right, factor loadings (minmax normalized) are shown for every neighborhood in the treated condition. Each neighborhood was assigned to the cluster that most frequently appears in the corresponding cells (by majority voting). Row color on the right: cluster assignment of a neighborhood. Neighborhoods are grouped by their assigned cluster and ordered by their average score for the corresponding factor. Only annotated clusters are shown. Rows in heatmaps are in direct correspondence. I.e., for every neighborhood on the right, in the corresponding row on the left the pre-images of this neighborhood were derived from the optimal transport solution 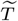 and their corresponding weighted average loading profile is shown. Row color on the left: most likely cluster assignment in the pre-image. To reflect the density of cells in each neighborhood, the thickness of a row corresponding to a neighborhood is proportional to its density in the treated condition (both left and right). Top gene markers for the shared component *W* for the DA factor are shown on the right. b) Condition-specific factors *V*_*c*_ contain information about local transcriptomic changes in cell states that are common between untreated (U) and treated (T) conditions. *V*_*T*_, the condition-specific factor for the treated condition, indicates genes that are upregulated locally in the psMEK treatment condition. *V*_*U*_, the wild type-specific factor for the untreated condition, indicates genes that are downregulated locally in the psMEK treatment condition. c) *V*_*T*_ for the DA factor indicates upregulated genes in dorsal animal cells in the treated condition. Loadings for top 30 genes are shown (and filtered for differential expression between conditions in the corresponding cells, with *>* 0.5 absolute log-foldchange and *<* 0.05 adjusted p-value cutoff). The underlined genes are an endothelial gene *she* and the dorsalizing factor *chrd*. d) *V*_*U*_ for the M factor indicates downregulated genes in marginal cells (filtered as in c)). Endodermal genes *lft2, sox32, gata5* and *gata6* are underlined.

Concurrently, there was expansion of the AL, SL and EL compartments at the expense of the VA compartment through over-expression of the corresponding factors. Interestingly, the increase in DA factor expression extends into the SL and EL state, generating highly aberrant states that express a mixture of new and wild type transcriptional signatures. We used the *V*_*c*_ components of DAISEE, which report local transcriptomic changes as treatment-specific DAISEE factors, to show that transcriptional program mixing is apparent even in cells that are common across conditions and are not entirely reprogrammed. The genes listed in the treatment-specific *V*_*c*_ component for a given factor indicate the upregulated genes upon psMEK activation in the corresponding cluster of cell states (Fig. 5b). E.g., the treatment-specific *V*_*c*_ component provided by DAISEE for the DA factor indicates that DA cells overexpress several genes upon psMEK treatment (Fig. 5c). One of these overexpressed genes is *chrd*, usually lowly expressed in wild type DA cells, with its expression more pronounced in D and MD cells (see Supp. Fig. 4, top MD factor genes). Alongside *chrd*, the gene *she*, also an endothelial gene usually expressed at later stages in the developing zebrafish vasculature, becomes overexpressed in DA cells upon psMEK treatment.

So far, our analysis primarily found changes in cell states outside of the margin. However, we expected to find some transcriptional responses in the margin, where Nodal signaling regulates FGF-driven ERK activation to ensure balanced specification of mesoderm and endoderm precursors (17). Endoderm induction is a stochastic event that is negatively modulated by FGF within a time window of Nodal signaling (18). Since psMEK treatment both amplifies and extends ERK activity in the embryo, we had expected psMEK treatment to prevent bipotential mesendoderm progenitors from switching into the endodermal fate. It is possible that negative feedback, known to buffer ERK activation when ectopic signals are present alongside ligand inputs, could occur where cells receive both ligand-derived and ectopic ERK inputs, and tamper ectopic psMEK-driven ERK signals in the margin (23). We nevertheless searched for subtle transcriptional changes in the margin using the untreated condition-specific *V*_*c*_ components of the M factor, pointing to down-regulated genes (Fig. 5d) in the margin. We indeed found that in the marginal cells, markers for endoderm progenitors *sox32* and *lft2* become downregulated upon psMEK-treatment, along with endodermal genes *gata5* and *gata6*. This result suggests attenuation of endoderm specification in psMEK-activated embryos.

## Discussion

Signaling pathways are necessary cues that drive uncommitted cells towards specific transcriptional states in a developing embryo. Developmental signals are often transient, whereas deregulated, sustained signal transduction in embryogenesis can lead to disease (44). Understanding the multifaceted changes in heterogeneous gene expression states in embryos experiencing spurious signaling requires single-cell approaches. However, single-cell studies have only perturbed signaling by removing pathway components (6, 45) which potentially only decreases the cell state diversity. Here, we find evidence that amplifying signaling can generate new cell states.

High-dimensional heterogeneous single-cell datasets like the ones we collected often require data integration algorithms in order to interpret. Data integration algorithms can provide an accurate comparison of common states across conditions, and identification of novel states that appear upon perturbation. Some available algorithms do not correct for confounding factors such as batch effects while others do not provide an easily interpretable framework, e.g., through factor analysis provided by iNMF methods. We presented DAISEE, an algorithm that combines the advantages of previously available iNMF-based approaches with explicit batch effect correction. DAISEE enabled construction of an ambient space in which cell state shifts and local, condition-driven cell state responses can be characterized (Fig. 6). DAISEE promises to be broadly applicable to transcriptional measurements with scRNA-seq and multiplexed RNA imaging, as well as single-cell measurements of other aspects of the cell state like chromatin accessibility, epigenetic profile, or genome organization (46, 47).

**Figure 6.**
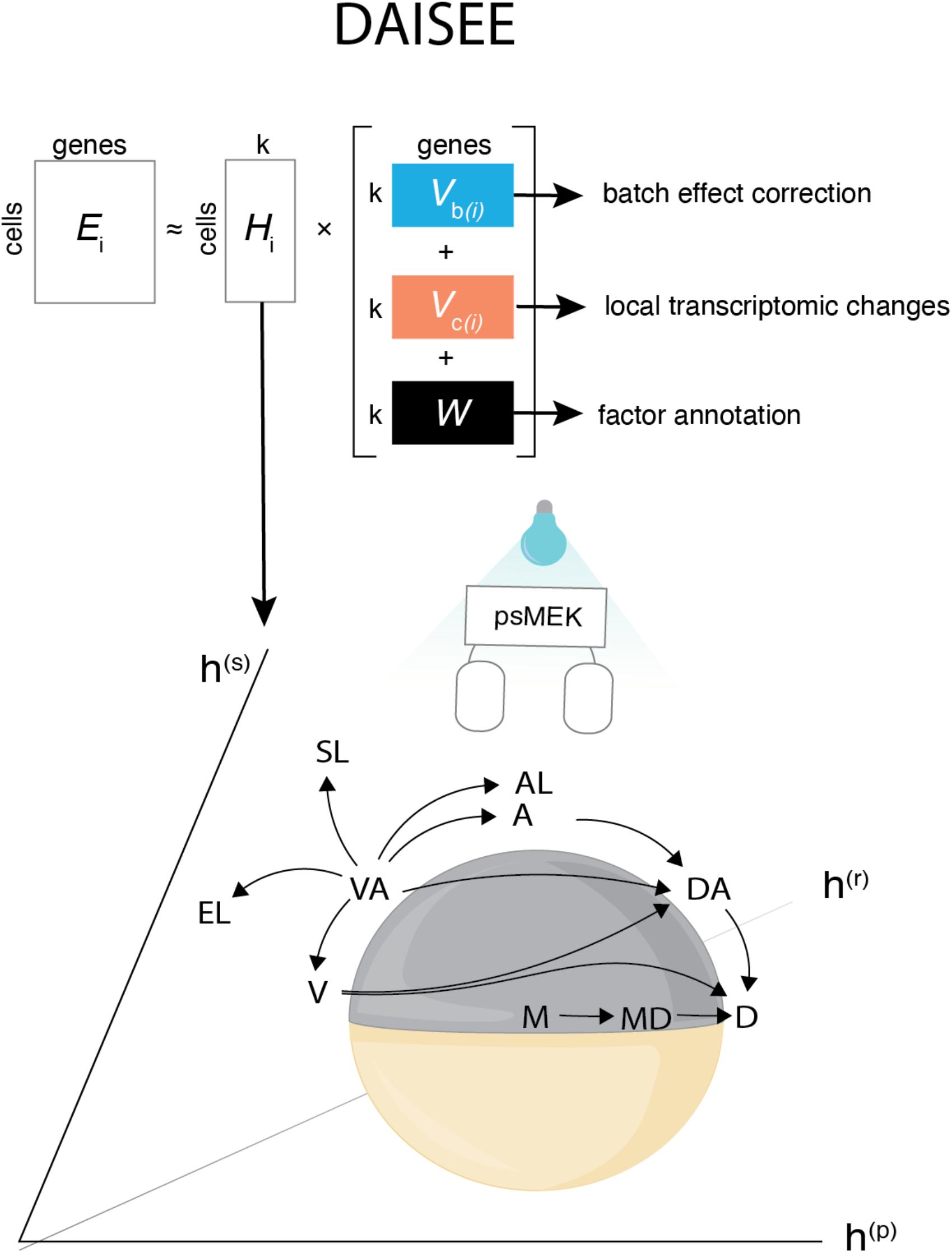
Summary of DAISEE. DAISEE is a framework for integrating high-dimensional single-cell data from multiple experimental conditions. It contains components that allow for batch correction (*V*_*b*_(*i*)) and data annotation (*W*). Cell factor scores (*H*_*i*_) are used for low-dimensional embedding that allows to study redistribution of cell states and discover new states. The *V*_*c*_(*i*) component describes local transcriptomic changes. We identified how psMEK treatment reassigned cell states in the zebrafish 50% epiboly embryo (arrows).

In this study, we used DAISEE to map transcriptional state shifts in 5 hour old zebrafish embryos experiencing hyperactive ERK signaling, and discovered several simultaneous directions of change including dramatic deviations from the normal transcriptomic landscape of the early embryo towards endothelial-like states (Fig. 6). We expect that mechanistic dissection of this state will shed light on the origins of abnormal blood vessel development, which is common in a broad class of developmental diseases that are caused by hyperactivity of the ERK pathway (44, 48). In later stages of zebrafish development, FGF ligands repress rather than promote endothelial fate (49). It is therefore unclear whether endothelial genes could be regulated by FGF-responsive genetic elements. Interestingly, oncogenic ERK pathway-activating mutations in somatic tissue also drive cerebral arteriovenous malformations likely by endothelial cell proliferation (50). Thus, upregulation of endothelial genes by ERK may require pathological levels of ERK signaling, and can be modeled and studied in the early zebrafish embryo with wide implications for human health. We suggest that dynamic measurements of cell state changes throughout early embryogenesis at multiple levels of gene regulation, from transcription factor binding, to chromatin accessibility and gene expression, will be crucial to address these open questions about ERK-mediated endothelial gene regulation.

## Code availability

DAISEE is written in python and is freely accessible at https://github.com/MariaAvdeeva/DAISEE

## Author Contributions

A.L.P. and M.A. designed the study and wrote the paper with contributions from S.Y.S. and R.D.B. A.L.P, V.G. and T.M. performed the experiments. M.A. developed and applied the computational data analysis tools.

## Acknowledgments

We thank the members of the S.Y.S. and R.D.B. laboratories and Flatiron Institute for helpful discussions. Sequencing was performed at the Lewis Sigler Institute Genomics Core Facility by Jennifer Miller with support of the director, Wei Wang. Imaging was performed with support from the Confocal Imaging Facility, a Nikon Center of Excellence, in the Department of Molecular Biology at Princeton University with assistance from Gary Laevsky. We thank Lucy Reading-Ikkanda for assistance with graphic design of figures.

## Methods

### ScRNA-seq data collection and preprocessing

Injection and optogenetics: *Danio rerio* embryos were raised according to standard zebrafish husbandry procedures. The pwt line was used to set up mating pairs. Synchronously fertilized embryos were collected and prepared for injection before they reached the 1-cell stage. psMEK mRNA was transcribed from the PCS2 psMEK E203K plasmid using the Ambion mMessage Machine SP6 kit. 50 pg of psMEK mRNA in a 500 pL droplet was injected into the yolks of each wild type embryo via a PV280 Pneumatic PicoPump (World Precision Instruments). Injected embryos were placed in glass petri dishes and stored in a 32ºC incubator in an aluminum foil lined box. A home built 505 nm LED board was placed on top of the box to flood the embryos in the petri dishes with light. For the batch-controlled experiments (batches 3 and 4, see Supp. Fig. 7a), un-injected embryos were placed under the 505 nm LED lights alongside injected embryos. All zebrafish procedures and experimental protocols in this study were conducted in accordance with the Princeton University Institutional Animal Care and Use Committee.

Dissociation: Embryos were removed from the light box when they reached the 50% epiboly stage and dechorionated. Dechorionation was performed by briefly softening chorions with pronase, washing out the pronase, and manually removing the chorion with forceps. 20-25 embryos per condition were deyolked manually in embryo medium and collected in a 1.5 ml Eppendorf tube. The deyolked embryos were resuspended in 200 µl embryo medium and dissociated by first flicking the tube 20 times then pipetting up and down 3 times. 800µl PBS with .1% BSA was added to the dissociated cells, and cells were spun down into a pellet at 300xg for 30 seconds. The supernatant was removed, being careful not to disturb the pellet, and the pellet was resuspended in 15-20µl PBS with .1% BSA. The concentration of cells was adjusted by diluting in PBS with .1% BSA as appropriate for downstream 10X Genomics scRNA-seq library preparation. In batch controlled experiments, samples from all conditions were prepared simultaneously.

Sequencing: Samples of dissociated cells were submitted to Princeton University Genomics Core facility for preparation and sequencing. Dissociated cells were processed with the Chromium Single Cell 3’ Assay. cDNA synthesis and library construction were carried out according to the manufacturer’s protocol. cDNA libraries were amplified with PCR and sequenced on the NovaSeq 6000 system. The resulting sequencing data were analyzed using the standard 10X Cellranger pipeline. Transcriptomic libraries were mapped to a zebrafish reference transcriptome built from the GRCz11 genome assembly and feature-barcode matrices were generated.

QC and preprocessing: After cellxgene count matrices were generated, standard pre-processing steps were applied as follows. First, data for all the samples were concatenated into one count matrix. Low quality cells were filtered by retaining only the cells with >1000 reads, >500 genes expressed and <5% mitochondrial counts. Genes were then filtered by retaining only the genes that are expressed in >10 cells and have >50 read counts. Outlier genes with >100000 read counts were removed. The concatenated data were clustered using Leiden clustering (50 PCs, 20 neighbors). We used the clustering results to filter the genes with >90% correlation of expression profiles over clusters with at least one gene from a pre-defined set of housekeeping genes (see (42) for the details). After this, the samples were separated and clustered separately with standard parameters (2000 highly variable genes with seurat_v3 method from the scanpy package, 20 neighbors, 50 PCs were used for each sample individually). In this step, we aimed to remove remaining low-quality and the EVL cells from every sample. For every sample, we manually removed clusters with low library sizes as low-quality cells (with thresholds differing for every sample due to differences in their library size distributions). EVL clusters could be clearly identified by differential expression analysis (Wilcoxon test) due to their overexpression of standard EVL markers such as *krt4, krt5* or *krt8*. This resulted in 17,276 cells after filtering. We note that some EVL cells were still identified in the filtered dataset during subsequent analysis.

### DAISEE algorithm

DAISEE solves an iNMF problem for a set of samples incorporating the experimental design into the objective function. More precisely, for *N* samples *X*_*i*_ collected with *C* conditions and *B* batches, with sample *i* corresponding to condition *c*(*i*) and batch *b*(*i*), we seek dataset-specific loadings *H*_*i*_ and matrices of common factors *W*, condition-specific factors *V*_*c*_ and batch-specific factors *V*_*b*_ which are a solution to the following optimization problem:

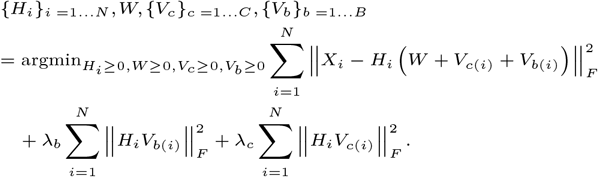

We approach this problem similarly to (1) by iterative updates of the loadings and factors through the following steps:

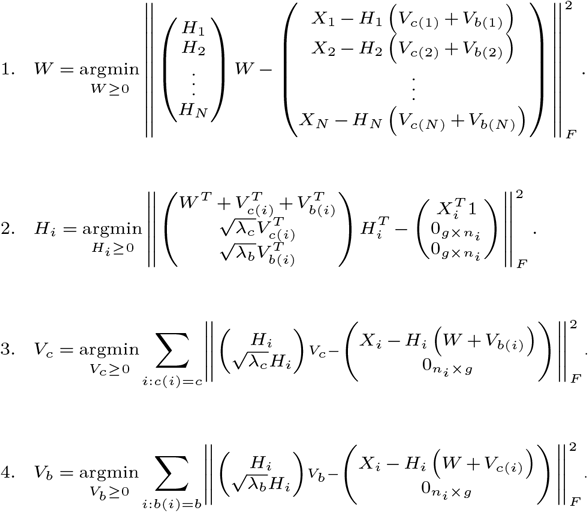

We solve each of the optimization problems independently using the block coordinate descent algorithm supplied by LIGER package (see (1) for details).

### Parameters for benchmarking experiments and details of implementation

For all benchmarking experiments (Supp. Figs. 1, 2), for LIGER, the default parameters of *k* = 30 and *λ* =5 were used. For DAISEE, we applied the same parameters, *k* = 30 and *λ*_*c*_ = 5, with varying *λ*_*b*_. The default 30 iterations of iNMF were applied for both.

Gene scaling was applied according to (1) at the sample level for both methods. Quantile normalization was applied to cell scores post-factorization for both methods. For the agreement metric, 100 nearest neighbors were used by default for jaccard similarity calculation for all benchmarks except the smaller simulated experiments where 15 nearest neighbors were used.

### Benchmarking DAISEE on scRNA-seq atlases

To demonstrate DAISEE on previously available scRNA-seq datasets, we chose two atlases from previous publications (25, 32). Immune atlas was aggregated for benchmarking purposes in (25) and is comprised of 96,348 human and mouse immune cells from different tissues that were collected over 23 samples by several laboratories. In this dataset, species were treated as perturbation for our purposes and we seeked to integrate the samples efficiently modeling the species-specific variability.

The mouse embryonic atlas (32) represents more homogeneous data and is comprised of 14,679 cells from mouse embryonic tissues collected over two consecutive stages, E7.0 and E7.5. In this dataset, we chose to model stage-specific differences in lieu of the perturbation. The data for both stages were collected over three batches resulting in a complex experimental design assumed by DAISEE.

### Applying DAISEE to the zebrafish embryo dataset

Top 1000 highly variable genes were used for all the analysis on the zebrafish embryo dataset. When applying DAISEE and LIGER on this dataset to compare their performance (Fig. 1), iNMF decomposition for both methods was run for 100 iterations and repeated 3 times, and the best replicate was chosen.

To finetune DAISEE on our newly collected data, we first followed the heuristic approach suggested in (1) to identify an appropriate number of factors for low-dimensional decomposition *k*. We calculate the Kullback-Leibler (KL) divergence (compared to uniform distribution) of the factor loadings for each cell and plot its median across cells as a function of *k*. The experiment for every *k* was repeated over 10 replicates. LIGER suggested that saturation in KL divergence would indicate that increasing the rank does not significantly change the sparsity of the factor loadings. We observed slow saturation of KL divergence (Supp. Fig. 3a) and chose to proceed with DAISEE at *k* = 30.

While we chose to use the default λ_*c*_ =5 parameter, we tuned λ_*b*_ parameter to achieve optimal alignment. Indeed, varying λ_*b*_ over a large domain on the log-scale, i.e., 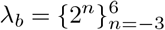 and repeating the experiment for every λ_*b*_ over 10 replicates, we discovered that the alignment saturates around λ_*b*_ =4 (Supp. Fig. 3b) which we fixed to proceed with the analysis.

After decomposing the zebrafish embryo dataset with 30 factors, we tested the factors for their batch and treatment specificity (Supp. Fig. 3d). More precisely, in accordance to (1), for every factor *j*, batch specificity was calculated as 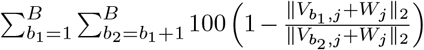.. Analogously, condition (treatment) specificity was calculated as 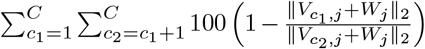. The threshold for batch specificity of factors was chosen at 6 to make sure that multiple factors exhibiting high treatment specificity and low batch specificity were included. Simultaneously, all but one of the 9 highly batch specific factors that were excluded demonstrated low treatment specificity (*<* 1.5). Excluding 9 batch-specific factors resulted in a perturbation to alignment and agreement of the DAISEE embedding setting alignment at 0.93 and agreement at 0.17.

For factor annotation, we compared our DAISEE factors to the previously published NMF factors from (6). The published factors were fitted to scRNA-seq data from wild type 50% epiboly zebrafish embryos, and were partially annotated. Top 30 markers for every identified component were published. To make a mapping, we intersected top 50 markers for DAISEE components corresponding to untreated condition (*W* + *V*_*U*_) with the published gene lists (Supp. Fig. 3c). As a result, we mapped all the previously annotated factors except the cell cycle component to one DAISEE factor with the largest intersection set (with the exception of MD for which two DAISEE components were retained).

### Simulation from iNMF framework

We used the DAISEE framework to simulate single-cell transcriptomics data for a complex experimental design. For these experiments, we chose the simplest possible design with two treatments collected across two batches (Supp. Fig. 1a). We reduced the parameter space by generating our toy single-cell data with *K* =3 factors only, and assuming *G* = 1000 as the number of genes considered, with *N* = 1000 cells in every sample. Every experiment consisted of 20 datasets (of 4 samples each) corresponding to a fixed pair (*α*_*c*_, *α*_*b*_) of condition-specific perturbation and batch effect strengths (see below for definitions). We allowed both *α*_*c*_ and *α*_*b*_ to run over the list [0.05, 0.2, 0.5, 1]. For every dataset, we simulated common (*W*), condition- (*V*_*c*_) and batch-specific (*V*_*b*_) iNMF factors. Furthermore, we simulated the corresponding cell scores *H* with an important simplifying assumption that condition can redistribute the cell scores in the dataset while batch does not. Finally, in every simulated sample, we varied the level of sparsity in our generated datasets by controlling the parameters of the library size distribution.

To be more precise, for every simulated dataset, we started by sampling the gene modules. Every 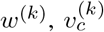 and 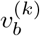 was sampled from a r distribution with shape 0.25 and scale 1 and size *G* = 1000. The shape of the distribution was fit to the biggest unperturbed sample (library-size normalized) in our zebrafish embryonic dataset and the scale parameter can be ignored due to downstream normalization. To control the size of the condition-specific and batch-specific effects, we normalized every sampled factor by their L1 norm (denoting the result by 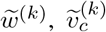 and 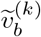. At a condition-specific perturbation strength parameter *–*_*c*_ and a batch effect parameter *–*_*b*_, for sample *s*, we considered 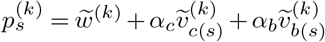 as the final gene module.

To sample condition-dependent cell scores 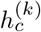 that would correspond to every sample *s* with *c*(*s*)= *c*, we chose a shape parameter 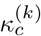 uniformly from [0.1, 1] and sampled *N* = 1000 cell scores from 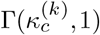. The cell scores were then normalized for every cell to sum up to 1. This resulted in a triangular geometry for every sample with condition-dependent concentration parameters (Supp. Fig. 1b).

Additionally, for every dataset (with 4 samples sharing sparsity level), a level of sparsity was chosen by controlling the mean of the library size distribution. After sampling a parameter *m* uniformly from [4, 7], library sizes for each dataset were sampled from a lognormal distribution with mean *m* and standard deviation 0.5 of the underlying normal distribution. This resulted in the datasets with sparsity varying from 45% to 94%, with the median sparsity of 80%.

After generating the gene loadings, cell scores and library sizes as specified above, for sample *s*, the RNA count for gene *j* of a cell *i* with library size *l*_*i*_ were sampled from a Poisson distribution with mean 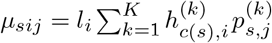.

Finally, when applying DAISEE to the simulated benchmarking experiments (Supp. Fig. 1c,d), a range of regularization parameters λ_*b*_ = [0, 0.001, 0.01, 0.1, 1, 10] were considered.

### Testing conditions for differential abundance

To study differential abundance between the conditions in the zebrafish embryo dataset, we chose to apply MILO (32). We first applied MILO to form overlapping neighborhoods in the *k*-nearest neighbor graph in the reduced, batch-corrected DAISEE space (dim = 21), choosing *k* = 100. To define the neighborhoods, MILO uses a refined sampling scheme initialized by sampling a fixed proportion of cell *p*. We chose *p* = 0.1 for our analysis which resulted in ≈100-300 cells per neighborhood. The strength of MILO is in its ability to test for differential abundance in overlapping neighborhoods. The method applies a negative binomial generalized linear model framework and uses a weighted FDR procedure to account for neighborhood overlap. We applied MILO with design=“ ∼condition” and applied FDR<=5% cutoff to the per-neighborhood hypothesis testing results (Supp. Fig. 5a). We refer the reader to the original MILO publication (32) for additional details.

### Optimal transport of cell state probability distributions between control and treatment

To study the transitions between control and treatment conditions in our data that resulted in the re-distribution predicted by MILO, we formulated an optimal transport problem. More precisely, MILO allowed us to extract the distributions of cells over neighborhoods in the control and treated conditions, with the corresponding probability distributions denoted by *µ* and *ν* respectfully. For any pre-defined cost-function *c*(*·, ·*) defined over pairs of neighborhoods, the optimal transport problem is searching for an optimal transportation map *T* : *X → X* transforming *µ* into ν and simultaneously minimizing the overall cost of transport, i.e.

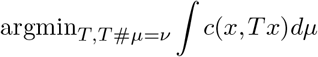

where # denotes the push-forward of measure. In practice, this problem (formulated by Gaspard Monge in 1781) is difficult to solve and is often substituted with a relaxation proposed by Leonid Kantorovich. The deterministic transportation map *T* is substituted by a *transportation plan*, i.e., a joint probability density π, with the problem becoming

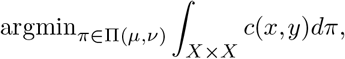

where Π (*µ, ν*), denotes the set of probability distributions on *X* × *X* with *µ* and *ν* as their corresponding marginals. Entropic regularization makes the problem strictly convex ensuring solution uniqueness and provable convergence. More precisely, a regularization term is added to the objective function as follows:

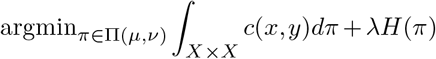

with the same conditions on π as above and *H*(π) denoting the entropy on the probability distribution π. We employed the implementation of entropic regularized optimal transport from the python pyopt package specifically devoted to optimal transport problems. To solve the optimization problem, we made use of the implementation of the Sinkhorn-Knopp algorithm within the package (`ot.sinkhorn`) and applied a small regularization parameter λ = 1*e* ≠ 3.

To define a cost function for transport between neighborhoods, we reasoned that it should reflect the transcriptomic distance between the neighborhoods in the batch-corrected space. More precisely, for every sample *E*_*i*_, the DAISEE algorithm provides a decomposition into low-dimensional (rank 21) matrices : *E*_*i*_ = *H*_*i*_(*W* + *V*_*c*(*i*)_ + *V*_*b*(*i*)_). To define the transcriptomic distance, we chose to correct for batch-specific effects but preserve the treatment-specific transcriptomic changes, i.e., we use Euclidean distance between the corrected transcriptomes 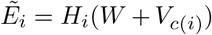, with quantile-normalized cell scores *H*.

Finally, every neighorhood gets assigned a state through majority voting of the corresponding cells. To define the cost of transport between 2 neighborhoods, *x* and *y*, we averaged all pairwise distances between the cells assigned to these neighborhoods. To calculate a transition probability from state *s*_1_ to state *s*_2_ upon treatment shown in Fig. 3, we use the optimal transportation plan 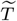 as follows:

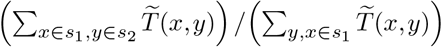

### Gene set enrichment analysis

We performed GSEA analysis for a collection of gene sets from a zebrafish embryonic development transcriptomic atlas (42). The atlas contains 194 gene sets of top differentially expressed genes in clusters of scRNA-seq data collected between 6 h.p.f. and 24 h.p.f. embryonic stages spanning from 11 clusters at 6 h.p.f. to 72 clusters of emerging tissues at 24 h.p.f. To run GSEA, we performed differential expression (DE) analysis between the clusters defined in Fig. 2b (Wilcoxon rank-sum test with Benjamini-Hochberg correction was applied on log-transformed library-size normalized data). For every cluster, we used the adjusted p-value (*pval_adj*) and the logfoldchange (*logFC*) of gene expression reported by DE results to rank the genes by ordering them by -log(*pval_adj*)*sign(*logFC*). To identify enrichment of a gene set at the top or bottom of a ranked list, GseaPreranked tool (43) was applied using the *conservative* classic scoring scheme. Only gene sets with at least 10 genes were used in this analysis. GSEA results were summarized in Fig. 4d and Supp. Fig. 8a.

### Hybridization Chain Reaction *in situ*

psMEK mRNA was transcribed from the PCS2 psMEK E203K plasmid using the Ambion mMessage Machine SP6 kit. 50 pg of psMEK mRNA in a 500 pL droplet was injected into each wild type embryo via a PV280 Pneumatic PicoPump (World Precision Instruments). Embryos were injected into one of the following areas: into the yolk at the one-cell stage, into the cell at the one-cell stage, or into one of two cells at the two-cell stage. Injected embryos and uninjected control embryos were immediately placed in 5 cm Pyrex glass dishes in an aluminum foil-lined box. The box was covered with a LED board, which emitted 505 nm light as previously described in (10). Embryos were incubated at 32°C and exposed to 505 nm light until 50% epiboly, at which point they were either dissociated for scRNA-seq or fixed in 4% paraformaldehyde overnight at 4°C for HCR. Fixed embryos were then washed in PBT, dechorionated, and gradually transitioned to 100% methanol for storage until the HCR protocol was started. WIK and Tu wild type strains were used.

The HCR RNA-FISH kit and probes for *fli1* and *lmo2* were purchased from Molecular Instruments. Molecular Instruments HCR RNA-FISH protocol for whole-mount zebrafish embryos and larvae (Danio rerio) version 8 was carried out with the following adjustments: for proteinase K treatment, embryos were treated with 1 mL of 100 µg/mL proteinase K for 5 minutes at room temperature; 3 µL of each probe was diluted into 200 µL of probe hybridization buffer for probe incubation; 6 µL of hairpin was diluted into 200 µL of amplification buffer for hairpin incubation; after hairpin solution removal, 1:500 Hoescht stain in 5X SSCT was added to sample tubes and incubated for 30 minutes at room temperature.

Embryos were deyolked and mounted in ProLong Diamond Antifade Mountant such that the vegetal pole touched the Fisherbrand microscope slide and the animal pole touched the VWR cover slip. Embryos were imaged using a Nikon A1 confocal microscope. Whole embryo images were obtained using the 10X objective with 20 µm Z-stacks. Cellular images were obtained using the 40X objective with 10 µm Z-stacks. pdDronpa from psMEK was detected with the 488 nm laser, Hoechst with the 408 nm laser, and HCR labeled RNA with the 647 nm laser. All images were obtained using the same laser settings.

### HCR spot quantification

Quantification of RNA spots detected in the HCR images was performed on individual z slices of the confocal stacks. Spots were detected using the Big-FISH python package. Nuclei were segmented using cellpose. For each image from an injected embryo spots and nuclei were counted only in regions of the image where the pdDronpa from the psMEK was detected. The same parameters for spot detection by Big-FISH were applied to all images. The number of spots normalized by the number of nuclei detected in the image was reported for each z slice.

