## Supplementary_figures for "Disrupted developmental signaling induces novel transcriptional states"

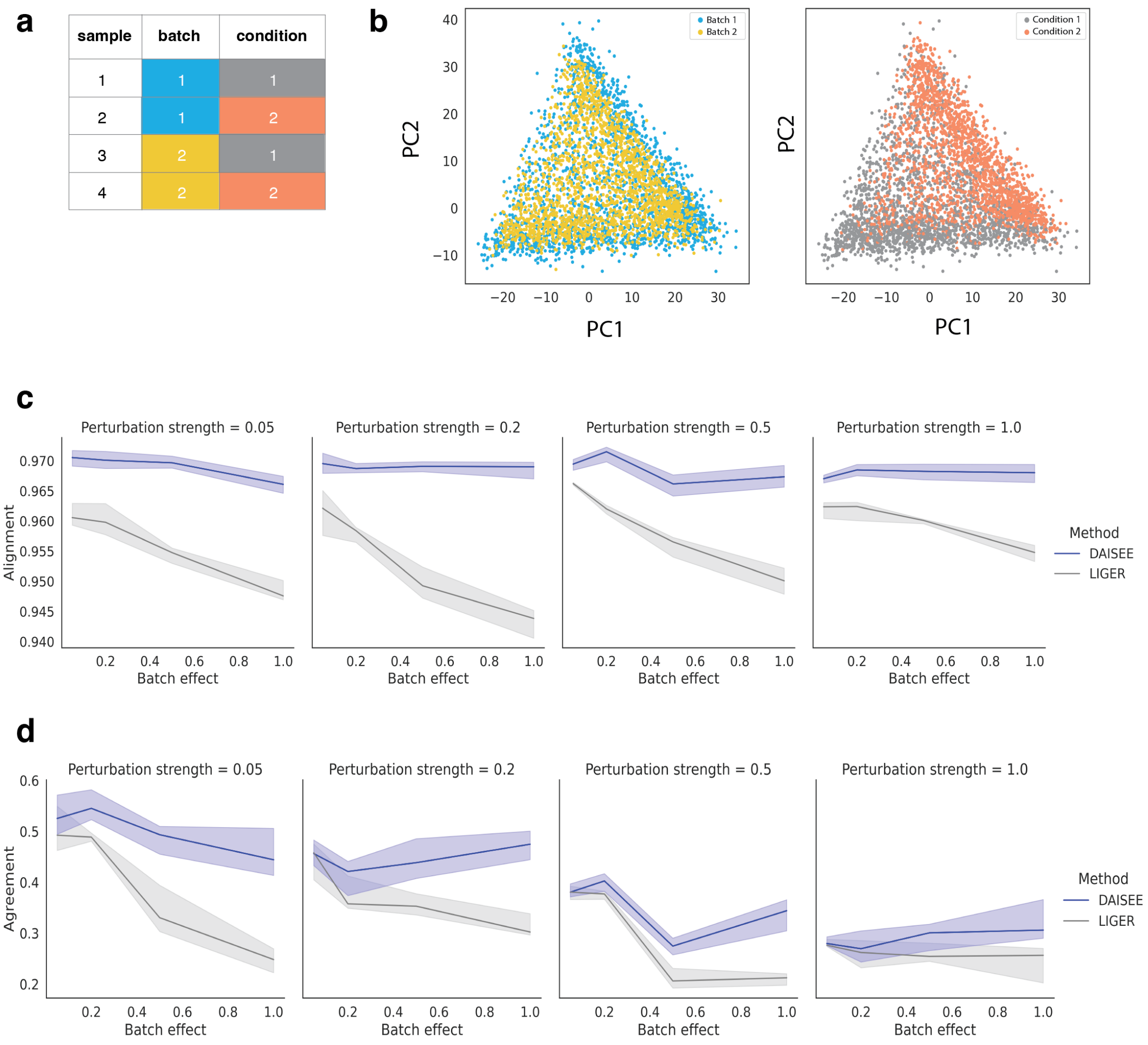


**Supplementary Figure 1. Benchmarking DAISEE vs LIGER on simulated single-cell data** a) A table showing the experimental design for every simulated dataset. b) Every simulated dataset consisted of 4 samples; an example with condition-specific perturbation strength $\alpha_c = 1$ and batch effect strength $\alpha_b = 1$ is shown. Principle component analysis (PCA) was applied to all cells in the library normalized samples; projection on first two PCs is shown. Batch-dependent differences (driven by $V_b$) only slightly perturb the samples after normalization (left), condition-dependent differences (driven not only by $V_c$ but also by $H$; right) are more apparent due to differences in cell score distribution via $H$. c) Alignment in simulated data. Every subfigure summarizes the results of simulated perturbation experiments, with every set of experiments featuring a different strength of perturbation $\alpha_c$ (Methods). $x$-axis: batch effect strength ($\alpha_b$). $y$-axis: batch alignment. Solid line: median DAISEE vs LIGER performance over 20 replicates per $\lambda_b$ value over a range of $\lambda_b$ values (Methods). 95% confidence interval is shaded. d) Same as c) with sample agreement on the $y$-axis.

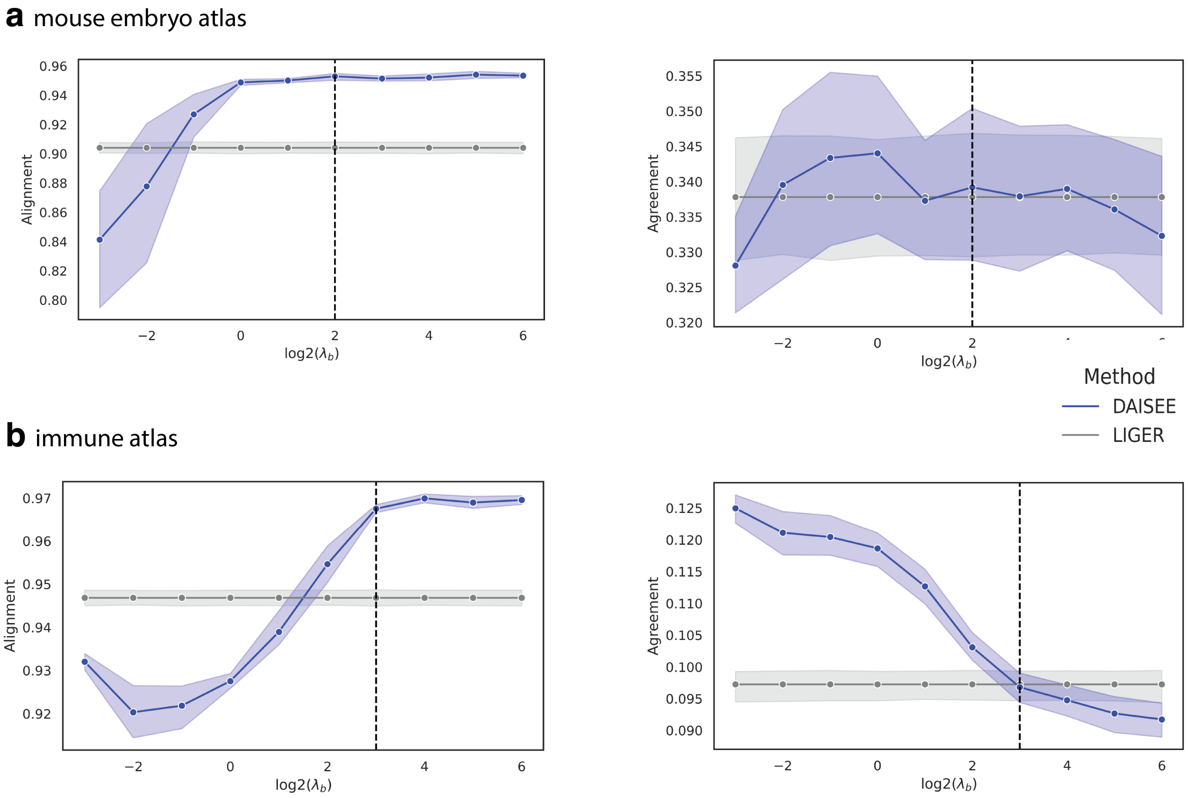


**Supplementary Figure 2. DAISEE outperforms LIGER on single-cell atlases** a) Benchmarks on a mouse embryo atlas (25). Blue: DAISEE, gray: LIGER. $x$-axis: batch-specific regularization parameter $\lambda_b$ from DAISEE, same results are shown for LIGER for every value since this parameter is absent in LIGER. $y$-axis: mean performance over 10 replicates is shown for both methods, the shading shows the 95% confidence interval. Left: alignment, right: agreement. Dashed line: an example of value of $\lambda_b$ that provides alignment improvement for similar agreement in DAISEE over LIGER. b) Same as a) for the immune atlas (32).

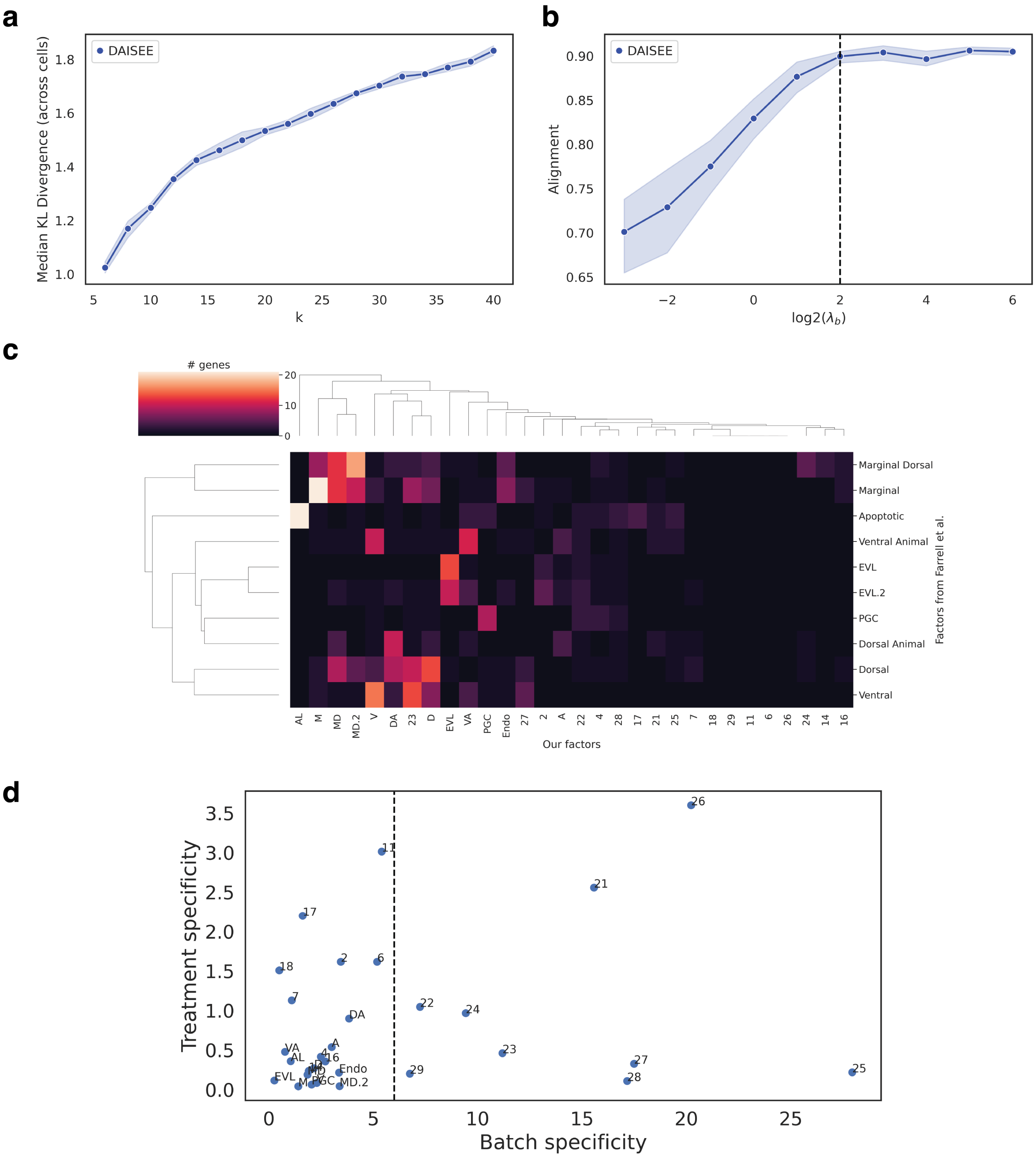


**Supplementary Figure 3. Applying DAISEE to the zebrafish embryonic dataset** a) $x$-axis: number of DAISEE factors $k$, $y$-axis: median Kullback-Leibler divergence (compared to uniform distribution) of the factor loadings over all cells (1). Mean and 95% confidence interval over 10 DAISEE experiments is shown. $k = 30$ was chosen. b) $x$-axis: Regularization parameter $\lambda_b$ (log2-scale), $y$-axis: alignment for a DAISEE experiment. Mean performance over 10 replicates, the shading shows the 95% confidence interval. Dashed line: $\lambda_b = 4$ was chosen. c) Clustermap of the size of the intersection of the sets of top 50 markers from DAISEE analysis with top 30 markers of the NMF factors derived from wild type 50% epiboly data in (6). d) Batch-vs-treatment specificity for each DAISEE factor fit to our zebrafish embryonic data (see Methods for more details). Dashed line: manually chosen cutoff on batch specificity that we applied.

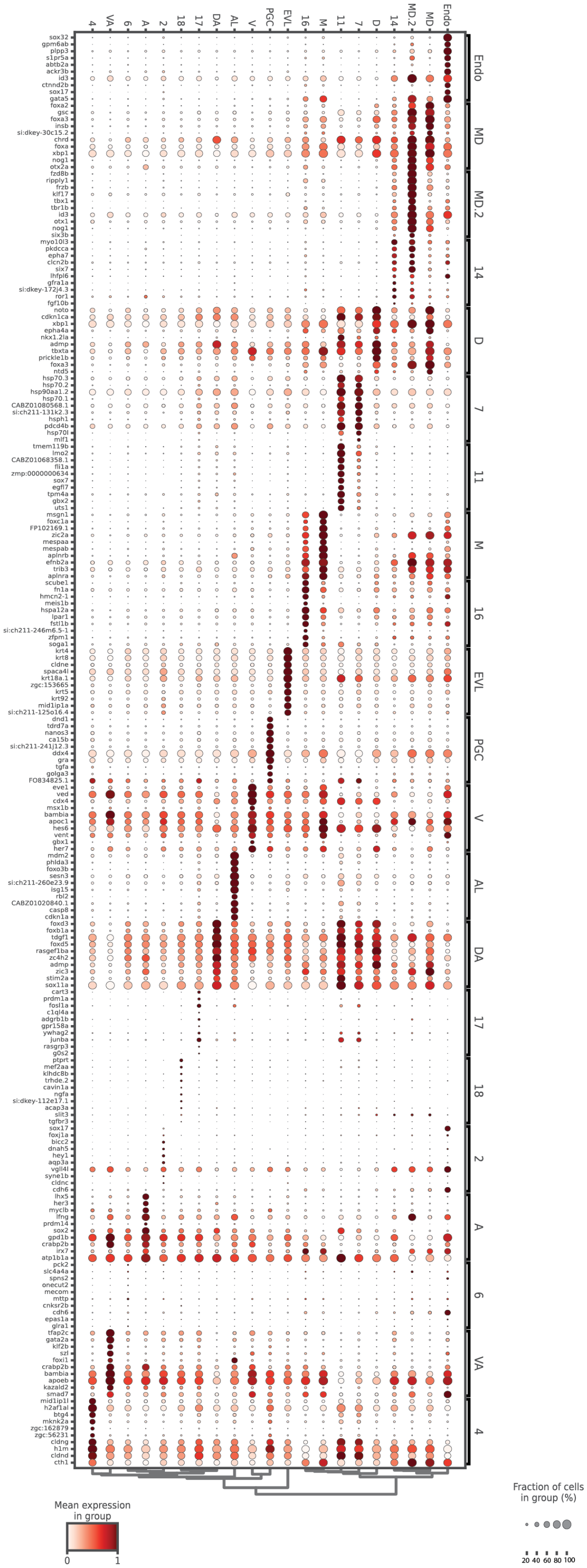

**Supplementary Figure 4. Top markers for DAISEE factors on the zebrafish embryonic dataset** Dotplot showing top expression of top 10 genes for the common component of each DAISEE factor, by cluster. Log-transformed library-size normalized expression is shown. The expression profile for every gene is then minmax normalized to vary between 0 and 1. Color of the dot shows mean expression per cluster, size of the dot corresponds to the fraction of cells expressing the gene.

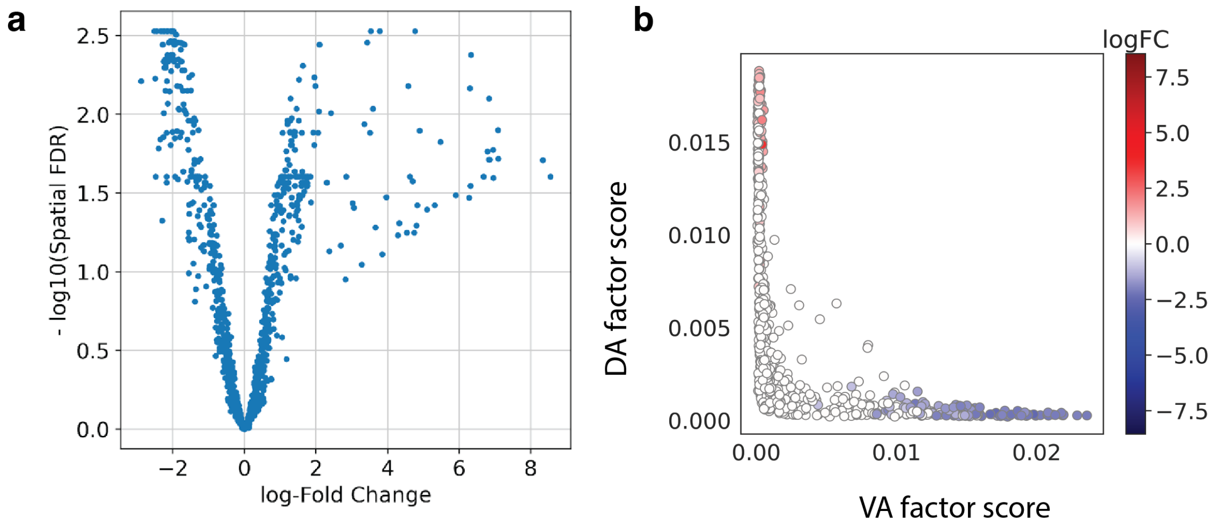


**Supplementary Figure 5. Details of the differential abundance analysis** a) Volcano plot for the differential abundance analysis. $x$-axis: logfoldchange of abundance between conditions, $y$-axis: spatial FDR calculated by MILO ($-log_{10}$ scale). $-log_{10}(0.05) \approx 1.3$ cutoff was chosen. b) $x$-axis: average VA factor loading in the neighborhood, $y$-axis: average DA factor loading in the neighborhood, color: same as in Fig.3.

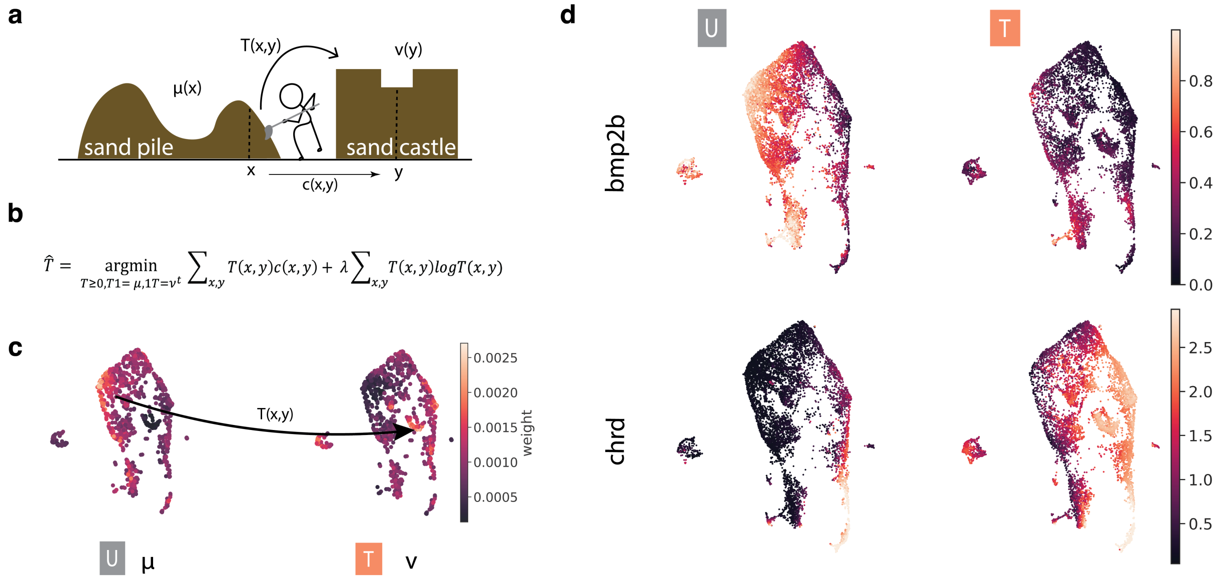


**Supplementary Figure 6. Applying optimal transport to the zebrafish embryonic dataset** a) A toy example of an optimal transport problem of transporting a pile of sand (represented by a probability distribution $\mu(x)$) into a sand castle (represented by a probability distribution $\nu(y)$). For a known cost function $c$, the optimal transportation map $T$ contructs the castle (i.e., transports $\mu$ into $\nu$) minimizing the total cost of transport. b) Optimization problem that we solved. The objective function includes entropic regularization for the transport map $T$, introducing an additional regularization parameter $\lambda$. c) In our data, we wish to solve an optimal transport problem that maps densities of cell states in the untreated condition to densities of cell states in the treated condition. d) UMAPs showing expression of *chrd* and *bmp2b* in the untreated and treated datasets (imputed using 50 nearest neighbors).

**a** light exposure among untreated condition samples

| sample | batch | condition | light |
|---|---|---|---|
| 1 | 1 | untreated (U) | dark |
| 2 | 3 | U | 500 nm |
| 3 | 4 | U | 500 nm |

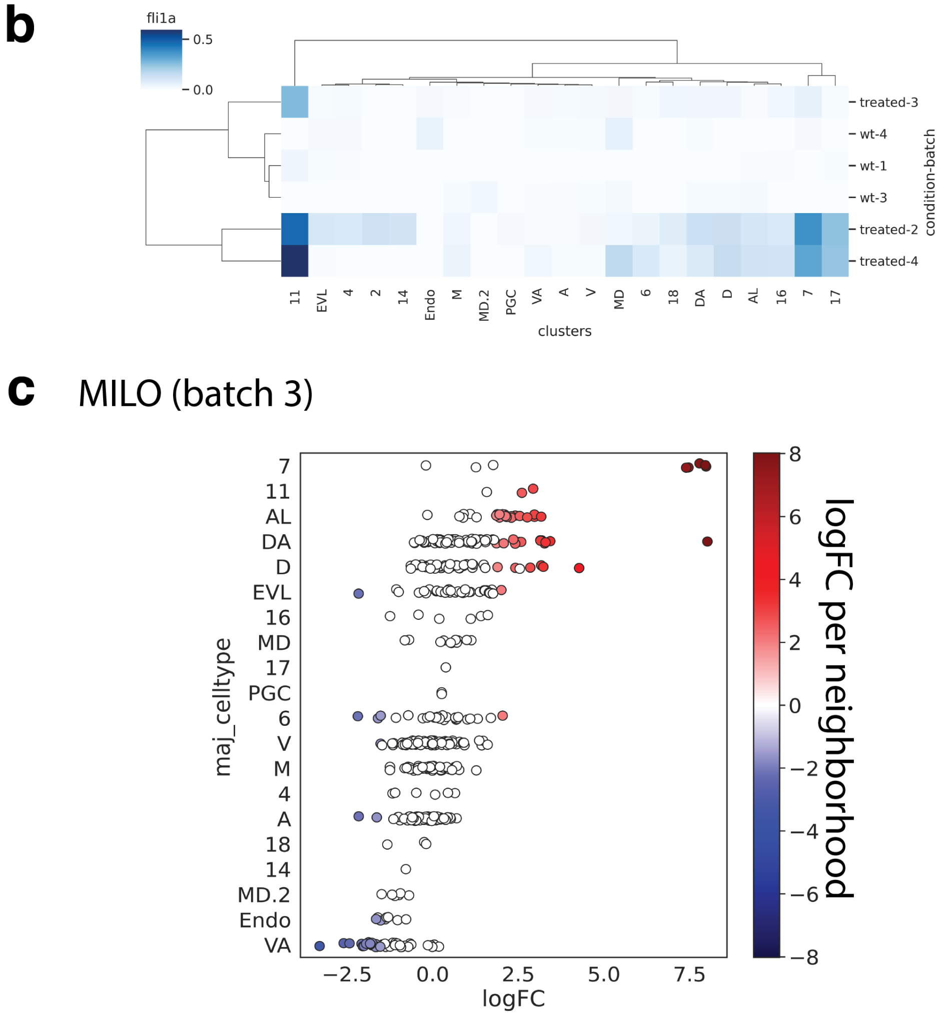


**Supplementary Figure 7. Novel states are observed in every replicate** a) A table showing light exposure among untreated samples. b) A clustermap showing average log-transformed library-size normalized expression of *fli1a*, one of the markers of Factor 11, per cluster of the zebrafish embryo dataset. c) Differential abundance testing results (as in Fig. 3a) applied to 3 samples corresponding to batch 3 only.

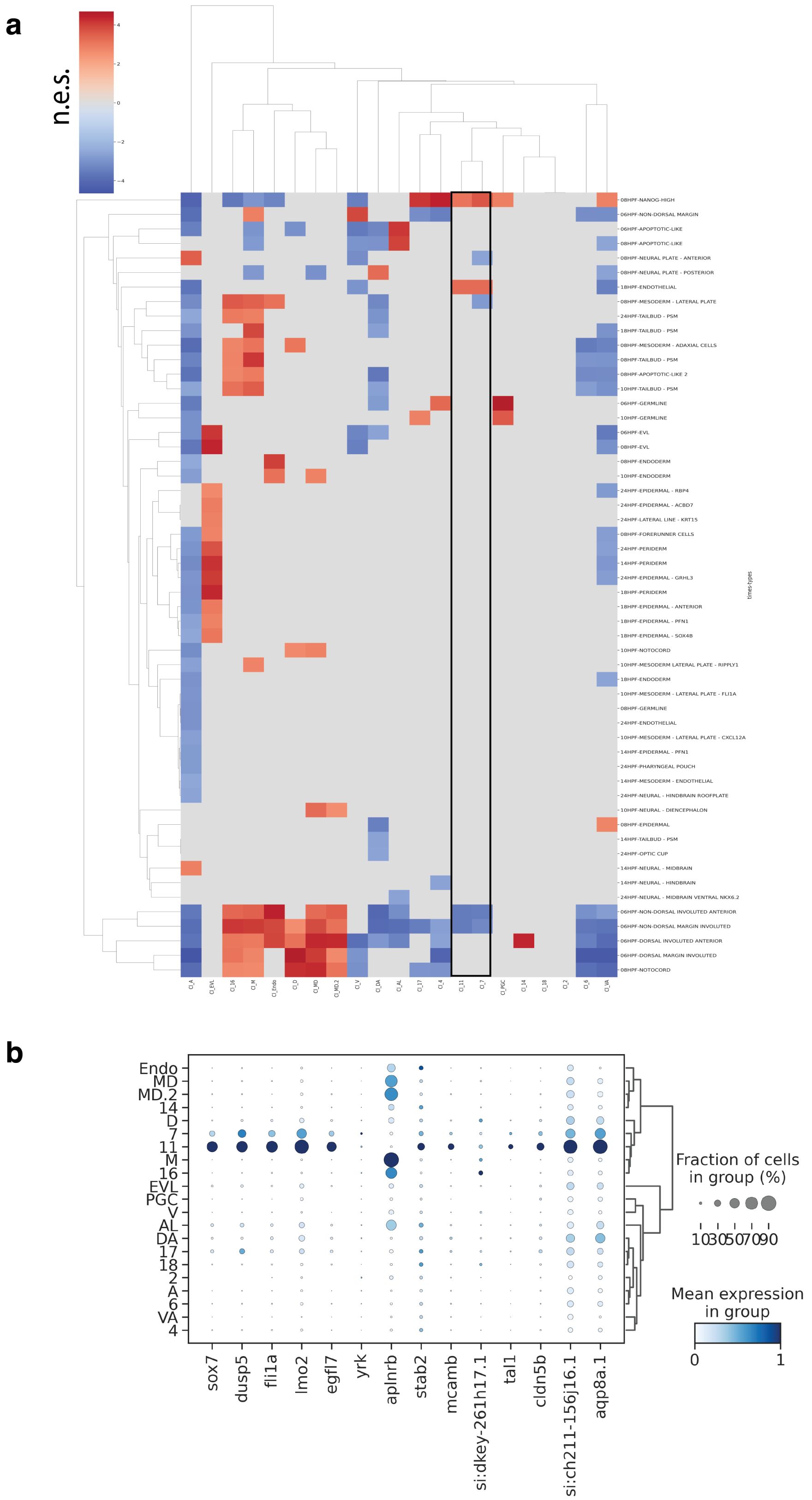


**Supplementary Figure 8. Global GSEA results and expression of endothelial transcriptional program genes in the zebrafish embryonic dataset** a) Normalized enrichment score after performing GSEA for all developmental time points. 18 h.p.f. endothelial genes are overexpressed in the 7 and 11 clusters of cell states (boxed). FDR q-value $< .001$ cutoff was applied. b) A dotplot showing log-transformed library-size normalized expression of the 18 h.p.f. endothelial markers in the clusters of the zebrafish embryo dataset. The expression profile for every gene is minmax normalized to vary between 0 and 1. Color of the dot shows mean expression per cluster, size of the dot corresponds to the fraction of cells expressing the gene.
